# Cancer-associated adipocytes mediate CD8^+^T cell dysfunction via FGF21-driven lipolysis

**DOI:** 10.1101/2024.10.18.618991

**Authors:** Sumiya Dalangood, Cegui Hu, Chenwei Yuan, Xiang Li, Wen Qiao, Hanjun Li, Rongyu Zhang, Luying Li, Peng Li, Xiang Yu, Wenjin Yin, Jinsong Lu, Jun Gui

**Author notes:** Correspondence to: **Dr. Jun Gui,** State Key Laboratory of Systems Medicine for Cancer, Renji-Med X Clinical Stem Cell Research Center, Ren Ji Hospital, School of Medicine, Shanghai Jiao Tong University, Shanghai, 200127, China **Email:**. These authors contributed equally (Sumiya Dalangood, Cegui Hu).

## Abstract

Cancer-associated adipocytes (CAAs) reprogram metabolic status of tumor microenvironment (TME). The metabolic crosstalk between CAAs and CD8^+^T cells in TME remains unclear. Here we report that CAAs undergo lipolysis, releasing free fatty acids that promote lipid peroxidation and disturb mitochondrial homeostasis in CD8^+^T cells, leading to their functional exhaustion. Importantly, we uncover that fibroblast growth factor 21 (FGF21) autocrinally drives CAA lipolysis through upregulating adipose triglyceride lipase (ATGL) via FGFR1/KLB-p38 signaling. FGF21 deletion in adipose tissue or ATGL inhibition impedes CAA lipolysis, mitigates lipid peroxidation, normalizes mitochondrial dynamics of CD8^+^T cells, and restores their effector function, consequently blunting tumor growth. Moreover, combining ATGL inhibitor with anti-PD-1 therapy synergistically enhances the antitumor activity of CD8^+^T cells, yielding greater therapeutic efficacy. Our findings highlight the pivotal role of CAA lipolysis in CD8^+^T cell dysfunction within TME, suggesting that targeting CAA lipolysis represents a valuable avenue for improving cancer immunotherapy.

## Introduction

Cancer immunotherapy using immune checkpoint inhibitors (ICIs) to enhance anti-tumor immune responses have achieved unprecedented success. Yet, only a small fraction of patients responds to ICIs due to the immunosuppressive tumor microenvironment (TME), highlighting the need to identify additional critical barriers that restrain antitumor immunity.^1,2^ TME is a highly heterogeneous ecosystem that contains tumor cells, matrix components, multiple infiltrating immune cells, and various stromal cells such as endothelial cells, fibroblasts, and adipocytes.^3^ Adipocytes, in particular, constitute a considerable portion of stroma in breast and colorectal tumors.^4^ Adipocytes surrounding the tumor site actively interact with tumor cells, leading to morphological and functional transformations.^5^ These adipocytes, termed as cancer-associated adipocytes (CAAs), exhibit dedifferentiated phenotypes and fibroblast-like characteristics.^6^ CAAs display delipidation to release lipid metabolites, such as free fatty acids (FFAs). Tumor cells utilize these FFAs as a source of fuel through fatty acid oxidation (FAO), facilitating their rapid growth.^7–9^ Besides, CAAs secret adipokines, inflammatory cytokines (e.g. IL-6, IL-1β) and chemokines (e.g. CCL-2, CCL-5) to support a pro-tumorigenic environment.^10–12^ To date, regarding the effect of CAAs on tumor progression, most studies have been focused on the crosstalk between CAAs and tumor cells. The role of CAAs in the regulation of tumor immune microenvironment remains largely unknown.

CD8^+^T cells are key effector cells in anti-tumor immune response, but they are often in a dysfunctional state with highly expression of immune inhibitory receptors such as PD-1, TIM-3, LAG-3.^13^ Increasing evidence suggests that metabolites and nutrients within TME reprogram CD8^+^T cell metabolism, thereby dictating anti-tumor immune response.^14,15^ Lipid accumulation is a common metabolic alteration in the TME.^16,17^ Accumulation of very-long-chain fatty acids or cholesterol induces tumor-infiltrating CD8^+^T cell exhaustion.^18,19^ Moreover, excessive lipid uptake promotes lipid peroxidation in CD8^+^T cells, impairing their anti-tumor function.^20,21^ However, the exact source of lipid abundance within TME is not very clear. Therefore, we intend to investigate whether CAAs contribute to the lipid enrichment, and how CAAs influence the lipid metabolism of CD8^+^T cells, thereby impacting their cytotoxic function.

Fibroblast growth factor 21 (FGF21) is a non-canonical member of fibroblast growth factor (FGF) superfamily.^22,23^ Physiologically, FGF21 is predominantly secreted from liver and adipose tissue, serving as an endocrine hormone. Under metabolic stress such as starvation or cold exposure, FGF21 is markedly induced to maintain energy homeostasis.^24,25^ In adipocytes, FGF21 acts as a cell-autonomous regulator via FGFR1-β-klotho (KLB) receptor complex to facilitate lipid disposal and induce thermogenesis.^26,27^ Our previous study demonstrates that tumor cells aberrantly secret FGF21 under TME stress, disrupting anti-tumor immunity by rewiring cholesterol metabolism of CD8^+^T cells.^28^ Here we find that CAAs secret higher levels of FGF21 when compared to normal adipocytes. FGF21 triggers CAA lipolysis by upregulating adipose triglyceride lipase (ATGL) via FGFR1/KLB-p38 signaling, releasing FFAs that promote lipid peroxidation and disturb mitochondrial homeostasis in CD8^+^T cells, consequently leading to their exhaustion. Adipocyte-specific deletion of FGF21 reverses ATGL-dependent lipolysis, reducing lipid uptake by CD8^+^T cells, normalizing mitochondrial dynamics, and restoring their anti-tumor immunity. Our findings underscore the significance of FGF21-ATGL-mediated CAA lipolysis in restricting CD8^+^T cell anti-tumor immune response, suggesting the therapeutic potential of targeting CAA lipolysis to boost anti-tumor immunity.

## Results

### CAAs undergo lipolysis and release free fatty acids

To understand the lipid metabolic characteristics of CAAs, we collected subcutaneous inguinal adipose tissue from wildtype (WT) C57BL/6 mice to obtain primary preadipocytes, which were differentiated into mature adipocytes. After that, the mature adipocytes were educated by breast cancer cells E0771 or colon adenocarcinoma cells MC38 using Transwell co-culture system to generate CAAs (Figure S1A). Non-tumor cell-educated WT adipose tissue adipocytes (AADs) served as control. We noticed that the morphology of adipocytes was changed after tumor cell education (Figure S1B). Meanwhile, the lipid content of these CAAs was significantly reduced when compared to the WT adipocytes as shown by Oil Red O staining and BODIPY-493 flow cytometry analysis (Figure 1A-B). We also harvested the adipose tissue at tumor-adjacent site from E0771 and MC38 tumor-bearing mice to obtain CAAs (Figure S1C). Adipocytes from WT tumor-free mice served as control. Similarly, tumor-adjacent CAAs exhibited much lower lipid content in contrast to WT AADs (Figure 1C-D). Indeed, the expression of marker genes for mature adipocytes was notably decreased in CAAs (Figure S1D), whereas the expression of proinflammatory cytokines and chemokines was greatly enhanced (Figure S1E). We further collected adjacent and distant adipose tissue from breast cancer patients who received total mastectomy. Accordingly, we observed that the morphology of tumor-adjacent adipocytes was shrink when compared to the normal distant adipocytes, along with markedly decreased lipid content (Figure 1E). These data suggest that CAAs exhibit delipidation.

**Figure 1.**
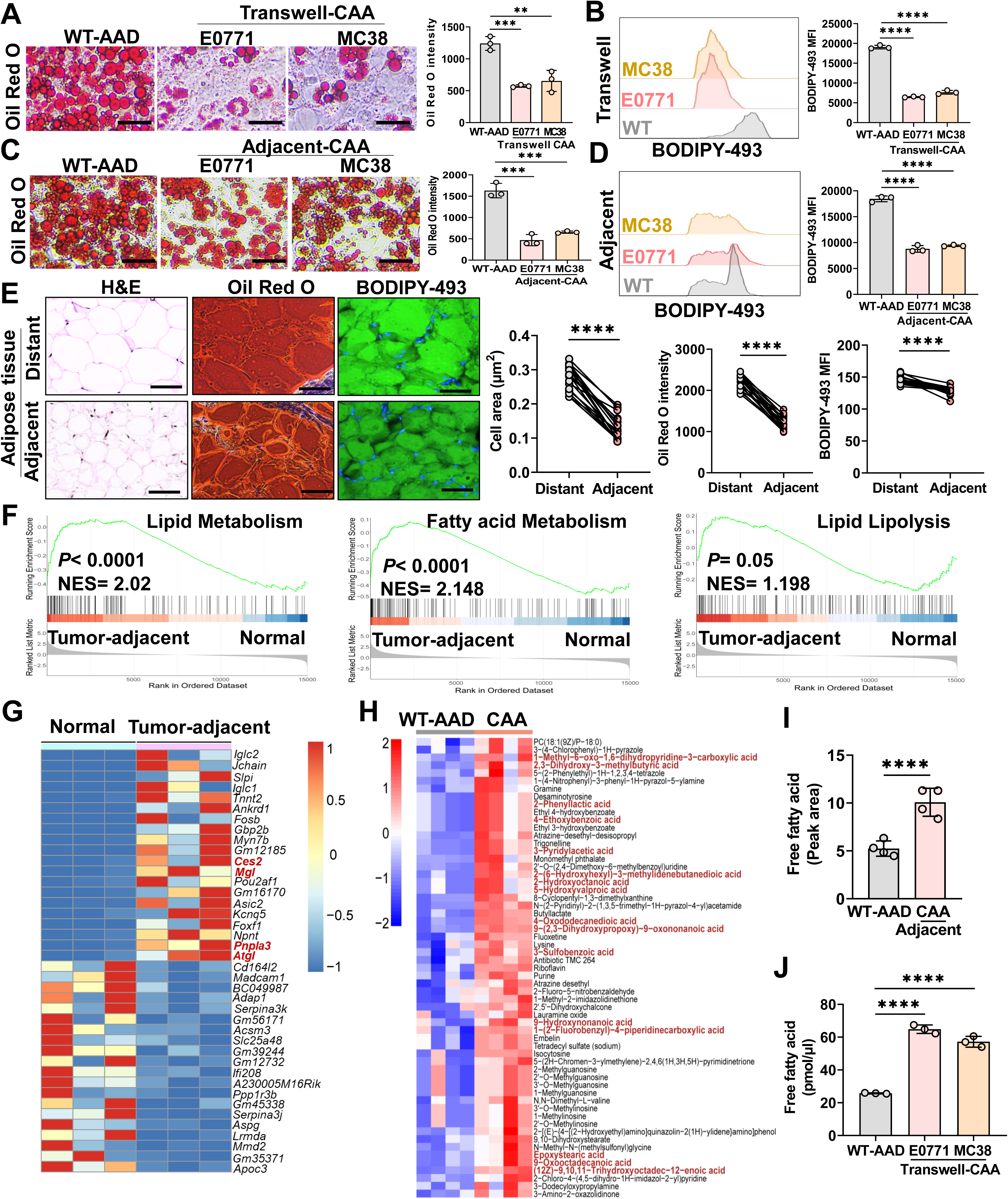
CAAs undergo lipolysis and release free fatty acids. **A.** Oil Red O staining of WT adipose tissue adipocytes (WT-AAD) and tumor cell (E0771 or MC38)-educated adipocytes (Transwell-CAA) (n = 3 biological replicates). **B.** Fluorescence-activated cell sorting (FACS) analysis of BODIPY-493 staining in WT AAD and tumor cell (E0771 or MC38)-educated adipocytes (Transwell-CAA) (n = 3 biological replicates). **C.** Oil Red O staining of WT-AAD or adipocytes from tumor-adjacent adipose tissue (Adjacent-CAA) (n = 3 biological replicates). **D.** FACS analysis of BODIPY-493 staining in WT-AAD or adipocytes from tumor-adjacent adipose tissue (Adjacent-CAA) (n = 3 biological replicates). **E.** Hematoxylin and eosin (H&E), Oil Red O and BODIPY-493 Immunofluorescence (IF) staining of tumor-adjacent and distant adipose tissue from breast cancer patients (n=18 patients). Scale bar: 100 μm. **F.** Gene set enrichment analysis (GSEA) of gene signatures of lipid metabolism, fatty acid metabolism, and lipolysis in WT normal adipose tissue and MC38 tumor-adjacent adipose tissue. **G.** Heatmap of differentially expressed genes in WT normal adipose tissue and MC38 tumor-adjacent adipose tissue. **H.** Liquid Chromatograph-Mass Spectrometer (LC-MS) analysis of metabolomics in the supernatant from WT-AAD or MC38 tumor-adjacent CAAs (n = 4 biological replicates). **I.** Metabolomics content of free fatty acids in the supernatant from WT-AAD or MC38 tumor-adjacent CAAs (n = 4 biological replicates). **J.** Quantification of free fatty acid levels in the supernatant from WT-AAD or tumor cell (E0771 or MC38)-educated adipocytes (Transwell-CAA) (n = 3 biological replicates). Data are presented as the mean ± SEM. \*\**P*<0.01, \*\*\**P*<0.001, \*\*\*\**P*<0.0001 by Student’s t test. (A, B, C, D, E, I, J).

To further clarify the features of CAAs, we performed RNA sequencing of MC38 tumor-adjacent adipose tissue. Gene set enrichment analysis (GSEA) revealed that genes associated with lipid metabolism, fatty acid metabolism and lipid lipolysis were significantly enriched in tumor-adjacent adipose tissue when compared to normal adipose tissue (Figure 1F). Specially, genes encoding lipid droplet-associate lipolytic enzymes, such as *Atgl*, *Pnpla3*, *Mgl* and *Ces2*, were upregulated in tumor-adjacent adipose tissue (Figure 1G). We validated that the expression of three rate-limiting lipases, along with proinflammatory cytokines, was markedly elevated in tumor-adjacent adipose tissue compared to distant adipose tissue from breast cancer patients (Figure S1F-G).

Triglycerides are the main energy molecule that is stored as lipid droplets within adipose tissues. Triglycerides can be hydrolyzed into glycerol and free fatty acids (FFAs) by those lipases.^29^ We performed the untargeted metabolomics analysis based on liquid chromatograph mass spectrometer (LC-MS) to determine the differential metabolites in the supernatant of MC38 tumor-adjacent CAAs. Of note, we found that more FFAs were released from CAAs (Figure 1H-I). By using FFA quantification assay, we validated that there was higher FFA content in the supernatant of CAAs (Figure1J). Collectively, all these data suggest that CAAs undergo lipolysis and release free fatty acids.

### CAAs promote fatty acid uptake and lipid peroxidation in CD8^+^T cells to induce their functional exhaustion

To investigate the impact of CAAs on lipid metabolism and effector function of CD8^+^T cells, we collected the conditioned media (CM) from WT AADs, transwell-CAAs and adjacent CAAs to treat CD8^+^T cells (Figure S2A). Lipid content in these cells was measured by BODIPY 493 staining at various time points following stimulation with anti-CD3/CD28. A progressive increase in neutral lipid content was observed in CD8^+^T cells during activation in response to CAA-conditioned media, with the most significant accumulation noted at 96 and 120 hr (Figure S2B). Subsequently, we analyzed the lipid uptake and peroxidation in CD8^+^T cells at 96 hr after anti-CD3/CD28 activation. Apparently, CD8^+^T cells exposed to CAA-conditioned media displayed higher lipid content, increased fatty acid uptake (BODIPY C16) and enhanced lipid peroxidation (BODIPY C11) when compared to those with WT AAD-conditioned media treatment (Figure 2A-B). Moreover, the expression of inhibitory receptors such as PD-1, LAG-3, and TIM-3 in CD8^+^T cells was notably elevated following CAA-conditioned media treatment (Figure 2C-D, Figure S2C-D). Conversely, the expression of cytotoxic functional molecules, including interferon (IFN) γ, TNF-α, Granzyme B, and Perforin, were markedly reduced in these CD8^+^T cells compared to the control group (Figure 2E-F, Figure S2E-F). All these data suggest that CAAs reprogram lipid metabolism of CD8^+^T cells, leading to their functional exhaustion.

**Figure 2.**
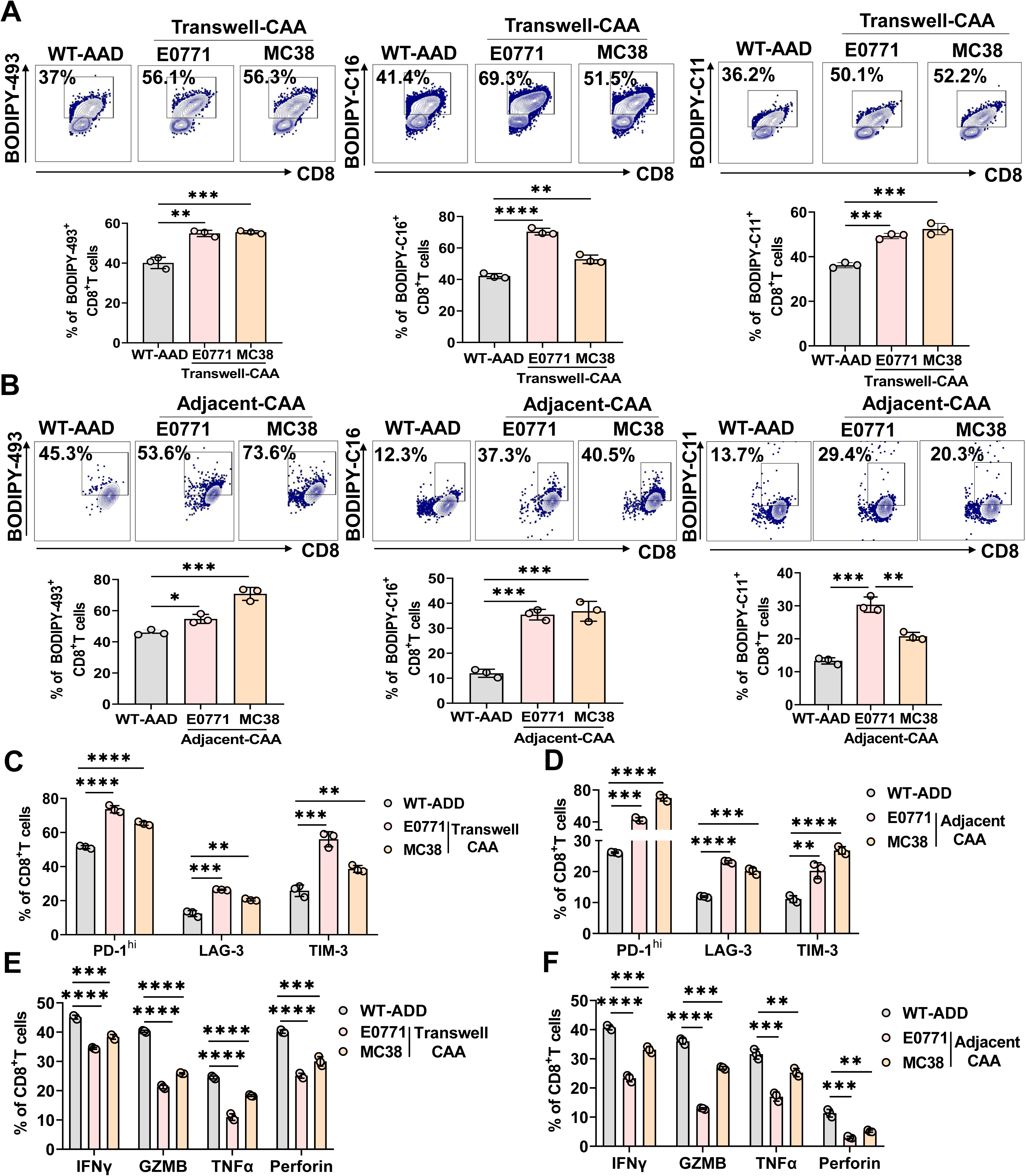
CAAs promote fatty acid uptake and lipid peroxidation in CD8^+^T cells to induce their functional exhaustion. **A-B.** FACS analysis of neutral lipid content (BODIPY-493), fatty acid uptake (BODIPY-C16) and lipid peroxidation (BODIPY-C11) in CD8^+^T cells treated with conditioned media from WT-AAD or tumor cell (E0771 or MC38)-educated adipocytes (Transwell-CAA) **(A)**, or E0771/MC38 tumor-adjacent CAAs **(B)** after activation by anti-CD3/CD28 for 96 hr (n = 3 biological replicates). **C-D.** FACS analysis of the percent of PD-1^hi^, LAG-3^+^, TIM-3^+^ of CD8^+^T cells treated with conditioned media from WT-AAD or tumor cell (E0771 or MC38)-educated adipocytes (Transwell-CAA) **(C)**, or E0771/MC38 tumor-adjacent CAAs **(D)** after activation by anti-CD3/CD28 for 96 hr (n = 3 biological replicates). **E-F.** FACS analysis of the percent of IFNγ^+^, Granzyme B^+^, TNFα^+^ and Perforin^+^ of CD8^+^T cells treated with conditioned media from WT-AAD or tumor cell (E0771 or MC38)-educated adipocytes (Transwell-CAA) **(E)**, or E0771/MC38 tumor-adjacent CAAs **(F)** after activation by anti-CD3/CD28 for 96 hr (n = 3 biological replicates). Data are presented as the mean ± SEM. \**P*<0.05, \*\**P*<0.01, \*\*\**P*<0.001, \*\*\*\**P*<0.0001 by Student’s t test (A, B) and two-way ANOVA (C, D, E, F).

### Selective deletion of FGF21 in adipocytes blunts tumor growth and restores the anti-tumor activity of CD8^+^T cells

In order to unravel the mechanism underlying CD8^+^T cell dysfunction induced by CAAs, we aimed to identify the key factor driving CAA lipolysis and FFA release. FGF21, an endocrine factor secreted by adipocytes, plays a crucial role in regulating lipid metabolism.^30^ Intriguingly, we observed a significant increase in secreted FGF21 levels in the supernatant of CAAs compared to WT AADs (Figure 3A), suggesting that FGF21 may contribute to CAA-mediated effects. To decipher the role of FGF21 in CAAs during tumor progression, we selectively ablated FGF21 in adipocytes by crossing *Fgf21*^fl/fl^ mice with *Adipoq*-Cre mice, generating *Fgf21*^ΔAD^ mice (Figure S3A). Mouse genotype in tail genomic DNA and Western blot confirmed the specific deletion of FGF21 in adipose tissue (Figure S3B-C). We then inoculated E0771 and MC38 tumor cells into *Fgf21*^fl/fl^ and *Fgf21*^ΔAD^ mice. Of note, the absence of FGF21 in adipocytes significantly suppressed tumor growth (Figure 3B and S3D). Moreover, the frequency of CD8^+^ tumor-infiltrating lymphocytes (TILs) was notably higher in *Fgf21*^ΔAD^ tumor-bearing mice (Figure 3C), whereas the percentages of dendritic cells (DCs), macrophages, polymorphonuclear myeloid-derived suppressor cells (PMN-MDSCs) and monocytic (M)-MDSCs in tumors were comparable between the two groups (Figure S3E).

**Figure 3.**
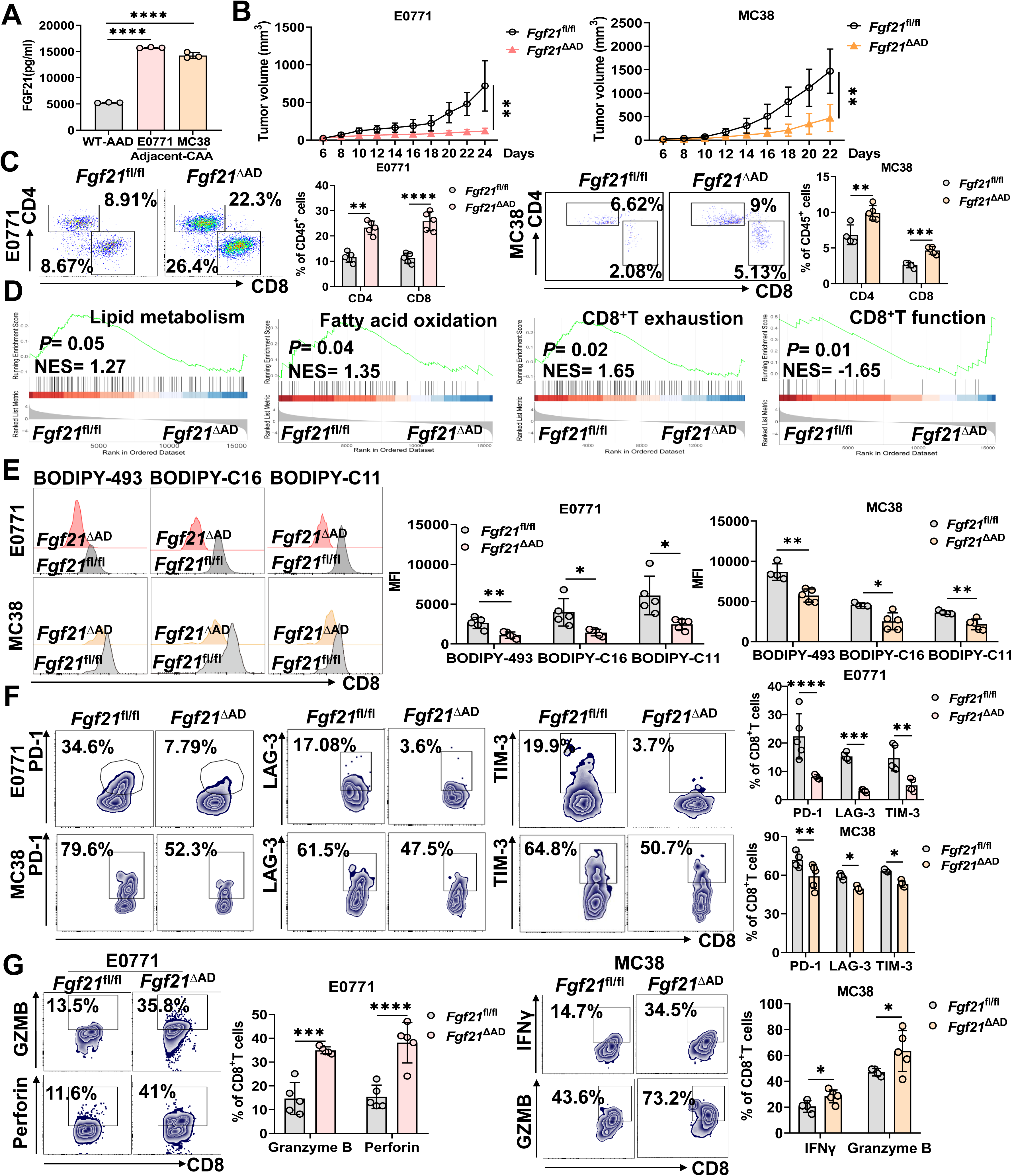
Selective deletion of FGF21 in adipocytes blunts tumor growth and restores the anti-tumor activity of CD8^+^T cells. **A.** ELISA analysis of FGF21 protein levels in the supernatant from WT-AAD and E0771/MC38 tumor-adjacent CAAs (n = 3 biological replicates). **B.** Tumor growth curve of E0771 and MC38 in *Fgf21*^fl/fl^ and *Fgf21*^ΔAD^ mice (n = 5 mice). **C.** FACS analysis of the percent of tumor-infiltrating CD4^+^ and CD8^+^T cells in E0771 and MC38 tumors from *Fgf21*^fl/fl^ and *Fgf21*^ΔAD^ mice (n =4-5 mice). **D.** GSEA analysis of gene signatures of lipid metabolism, fatty acid oxidation, CD8^+^T cell exhaustion and function in CD8^+^TILs sorted from MC38 tumors in *Fgf21*^fl/fl^ and *Fgf21*^ΔAD^ mice (n = 3 mice). **E.** FACS analysis of BODIPY-493, BODIPY-C16 and BODIPY-C11 staining in CD8^+^TILs of E0771 and MC38 tumors in *Fgf21*^fl/fl^ and *Fgf21*^ΔAD^ mice (n = 4-5 mice). **F.** FACS analysis of PD-1, LAG-3 and TIM-3 in CD8^+^TILs of E0771 and MC38 tumors in *Fgf21*^fl/fl^ and *Fgf21*^ΔAD^ mice (n = 4-5 mice). **G.** FACS analysis of IFNγ, Granzyme B, and Perforin in CD8^+^TILs of E0771 and MC38 tumor in *Fgf21*^fl/fl^ and *Fgf21*^ΔAD^ mice (n = 4-5 mice). Data are presented as the mean ± SEM. \**P*<0.05, \*\**P*<0.01, \*\*\**P*<0.001, \*\*\*\**P*< 0.0001 by Student’s t test (A) and two-way ANOVA (B, C, E, F, G).

To gain further insight into how adipocyte-derived FGF21 impact CD8^+^T cell activity, we sorted out CD8^+^TILs from MC38 tumor-bearing *Fgf21*^fl/fl^ and *Fgf21*^ΔAD^ mice, and performed RNA-sequencing (Figure S3F). GSEA revealed that pathways involved in lipid metabolism, fatty acid oxidation, and T cell exhaustion were significantly downregulated in CD8^+^TILs from *Fgf21*^ΔAD^ mice compared to controls. Conversely, genes associated with CD8^+^T cell functionality were upregulated in *Fgf21*^ΔAD^ mice (Figure 3D), indicating that FGF21 deletion in adipocytes may restore CD8^+^T cell activity. Indeed, we observed reduced lipid content, fatty acid uptake, and lipid peroxidation in CD8^+^TILs from *Fgf21*^ΔAD^ mice in contrast to *Fgf21*^fl/fl^ mice (Figure 3E). Accordingly, the proportion of exhausted CD8^+^TILs decreased (Figure 3F), while the expression of cytotoxic molecules in CD8^+^T cells increased (Figure 3G). Taken together, these results suggest that FGF21 specific deletion in adipocytes suppresses tumor growth and enhances the anti-tumor activity of CD8^+^T cells.

### CAAs inhibit CD8^+^T cell effector function in a FGF21-dependent manner

We proceeded to determine whether FGF21 directly contributes to the effects mediated by CAAs. We collected adipose tissues adjacent to E0771 and MC38 tumors from *Fgf21*^fl/fl^ and *Fgf21*^ΔAD^ tumor-bearing mice and differentiated into adipocytes, generating WT CAAs and FGF21-deficient CAAs. We found that the absence of FGF21 apparently restored lipid storage in CAAs, as evidenced by Oil Red O staining as well as BODIPY-493 flow cytometry analysis (Figure 4A-B). In line with this, the FFA levels were substantially lower in the supernatant of FGF21-deficient CAAs compared to the control CAAs (Figure S4A), indicating that FGF21 deficiency alleviated CAA lipolysis. We then treated CD8^+^T cells with the conditioned media from *Fgf21*^fl/fl^ and *Fgf21*^ΔAD^ CAAs. Notably, the lipid content, fatty acid uptake, and lipid peroxidation were all mitigated in CD8^+^T cells treated with conditioned media from FGF21-deficient CAAs (Figure 4C), as opposed to those treated with conditioned media from WT CAAs. Similarly, the proportion of exhausted T cells was significantly decreased following treatment with conditioned media from FGF21-deficient CAAs (Figure 4D and S4B). By the contrast, FGF21 ablation in CAAs led to the reinstatement of cytotoxic molecule expression in CD8^+^T cells exposed to CAA-conditioned media (Figure 4E and S4C). These data indicate that FGF21 is essential for CAA-mediated lipid peroxidation in CD8^+^T cells and the subsequent T cell dysfunction.

**Figure 4.**
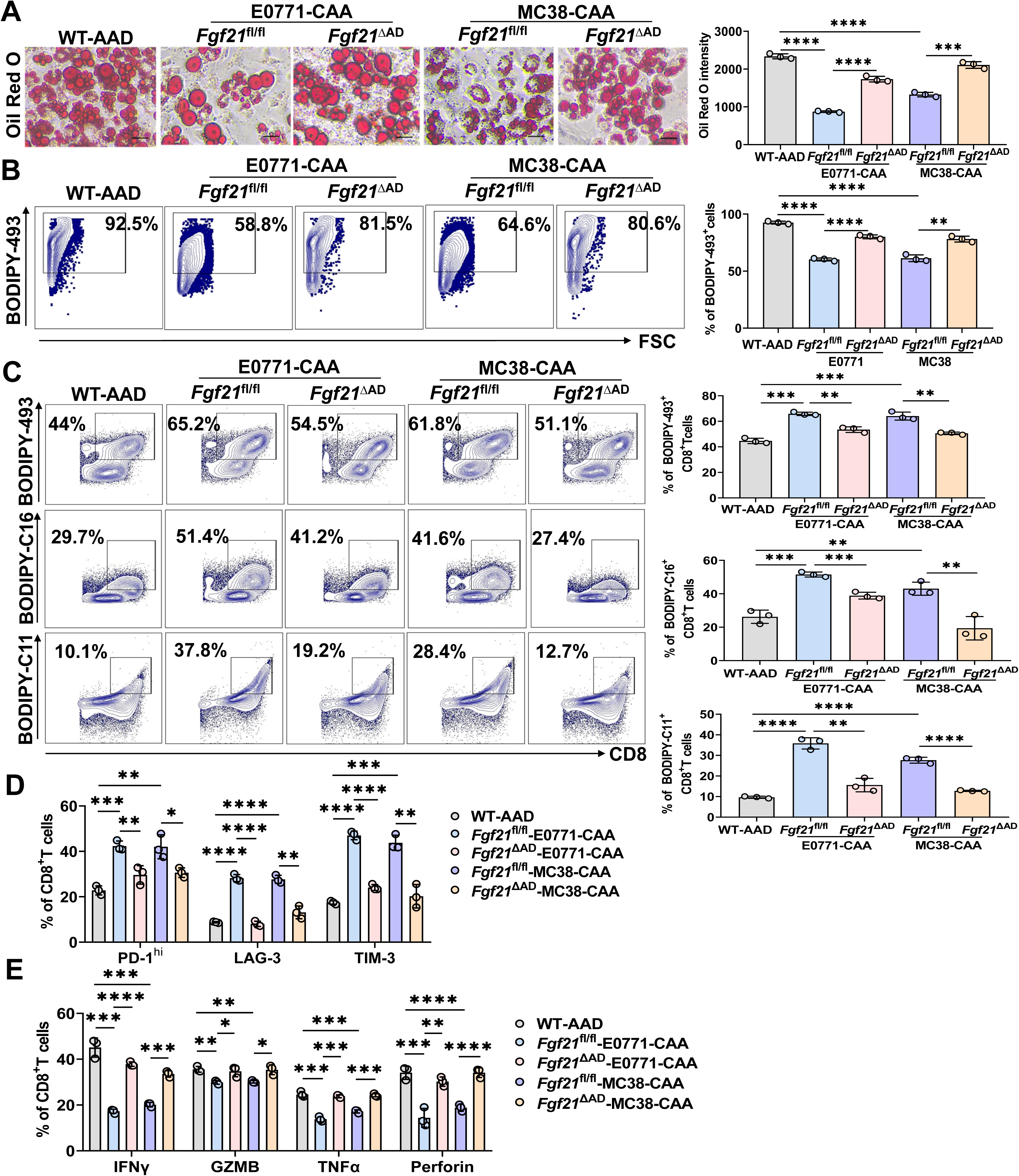
CAAs inhibit CD8^+^T cell effector function in a FGF21-dependent manner. **A.** Oil Red O staining of WT-AAD or E0771/MC38 tumor-adjacent CAAs from *Fgf21*^fl/fl^ and *Fgf21*^ΔAD^ mice (n = 3 biological replicates). **B.** FACS analysis of BODIPY-493 staining in WT-AAD or E0771/MC38 tumor-adjacent CAAs from *Fgf21*^fl/fl^ and *Fgf21*^ΔAD^ mice (n = 3 biological replicates). **C.** FACS analysis of BODIPY-493, BODIPY-C16 and BODIPY-C11 staining in CD8^+^T cells treated with conditioned media from WT-AAD or E0771/MC38 tumor-adjacent CAAs from *Fgf21*^fl/fl^ and *Fgf21*^ΔAD^ mice after activation by anti-CD3/CD28 for 96 hr (n = 3 biological replicates). **D.** FACS analysis of the percent of PD-1^hi^, LAG-3^+^ and TIM-3^+^ of CD8^+^T cells treated with conditioned media from WT-AAD or E0771/MC38 tumor-adjacent CAAs from *Fgf21*^fl/fl^ and *Fgf21*^ΔAD^ mice after activation by anti-CD3/CD28 for 96 hr (n = 3 biological replicates). **E.** FACS analysis of the percent of IFNγ^+^, Granzyme B^+^, TNFα^+^ and Perforin^+^ of CD8^+^ T cells treated with conditioned media from WT-AAD or E0771/MC38 tumor-adjacent CAAs from *Fgf21*^fl/fl^ and *Fgf21*^ΔAD^ mice after activation by anti-CD3/CD28 for 96 hr (n = 3 biological replicates). Data are presented as the mean ± SEM. \**P*<0.05, \*\**P*<0.01, \*\*\**P*<0.001, \*\*\*\**P*<0.0001 by Student’s t test (A, B, C) and two-way ANOVA (D, E).

### CAAs disturb mitochondrial homeostasis in exhausted CD8^+^T cells via FGF21

Given that excess FFAs undergo β-oxidation in Mitochondria, which are highly vulnerable to oxidative damage,^31^ we intended to assess whether CAA-mediated FFA uptake and lipid oxidation, triggered by FGF21, affect mitochondrial function in CD8^+^T cells. Analysis of RNA-seq data from CD8^+^ TILs revealed a significant downregulation of genes associated with mitochondrial dysfunction in CD8^+^TILs from *Fgf21*^ΔAD^ mice compared to *Fgf21*^fl/fl^ mice (Figure 5A). We then conducted Transmission Electron Microscope (TEM) to investigate mitochondrial morphology of CD8^+^T cells treated with CAA-conditioned media. Intriguingly, we observed increased mitochondrial fusion and elongation in CD8^+^T cells exposed to CAA-conditioned media, indicating that CAA-induced fatty acid oxidation disturbs mitochondrial homeostasis (Figure 5B). However, the phenomenon of mitochondrial fusion was diminished in CD8^+^T cells treated with media from FGF21-deficient CAAs (Figure 5B). Mitochondrial fusion is known to promote oxidative phosphorylation (OXPHOS) and reactive oxygen species (ROS) production, which in turn aggravates oxidative stress.^32,33^ We monitored the dynamic change of oxygen consumption rate (OCR) by Seahorse and ROS production by flow cytometry in CD8^+^T cells stimulated with anti-CD3/CD28 at different time points. Under normal condition, OCR was robustly elevated at 24 hr, but gradually declined by 96 hr (Figure S5A). Consistently, ROS production exhibited the similar tendency rising up at 24-48 hr, but declining by 72-96 hr. Notably, the proportion of ROS-high (ROS^hi^) CD8^+^T cells was minimal at 96 hr post-activation (Figure S5B). Previous data showed that CAAs promoted lipid accumulation and oxidation in CD8^+^T cells pronouncedly at 96 hr after stimulation (Figure S2B). In fact, both OCR and ROS^hi^ expression in CD8^+^T cells stimulating at 96 hr were markedly exaggerated following *Fgf21*^fl/fl^ CAA-conditioned media treatment when compared to the controls (Figure 5C-D). Yet, these parameters were normalized in cells treated with *Fgf21*^ΔAD^ CAA-conditioned media (Figure 5C-D). Additionally, ROS^hi^ CD8^+^T cells displayed much higher PD-1 expression compared to ROS^low^ CD8^+^T cells (Figure S5C), suggesting that ROS contributes to CD8^+^T cell exhaustion. Likewise, the proportion of ROS^hi^PD-1^hi^ CD8^+^T cells was greatly increased following *Fgf21*^fl/fl^ CAA-conditioned media treatment in contrast to the control treatment, while this enhancement was eliminated with *Fgf21*^ΔAD^ CAA-conditioned media treatment (Figure 5E). All these findings suggest that CAAs disrupt mitochondrial homeostasis in exhausted CD8^+^T cells via FGF21.

**Figure 5.**
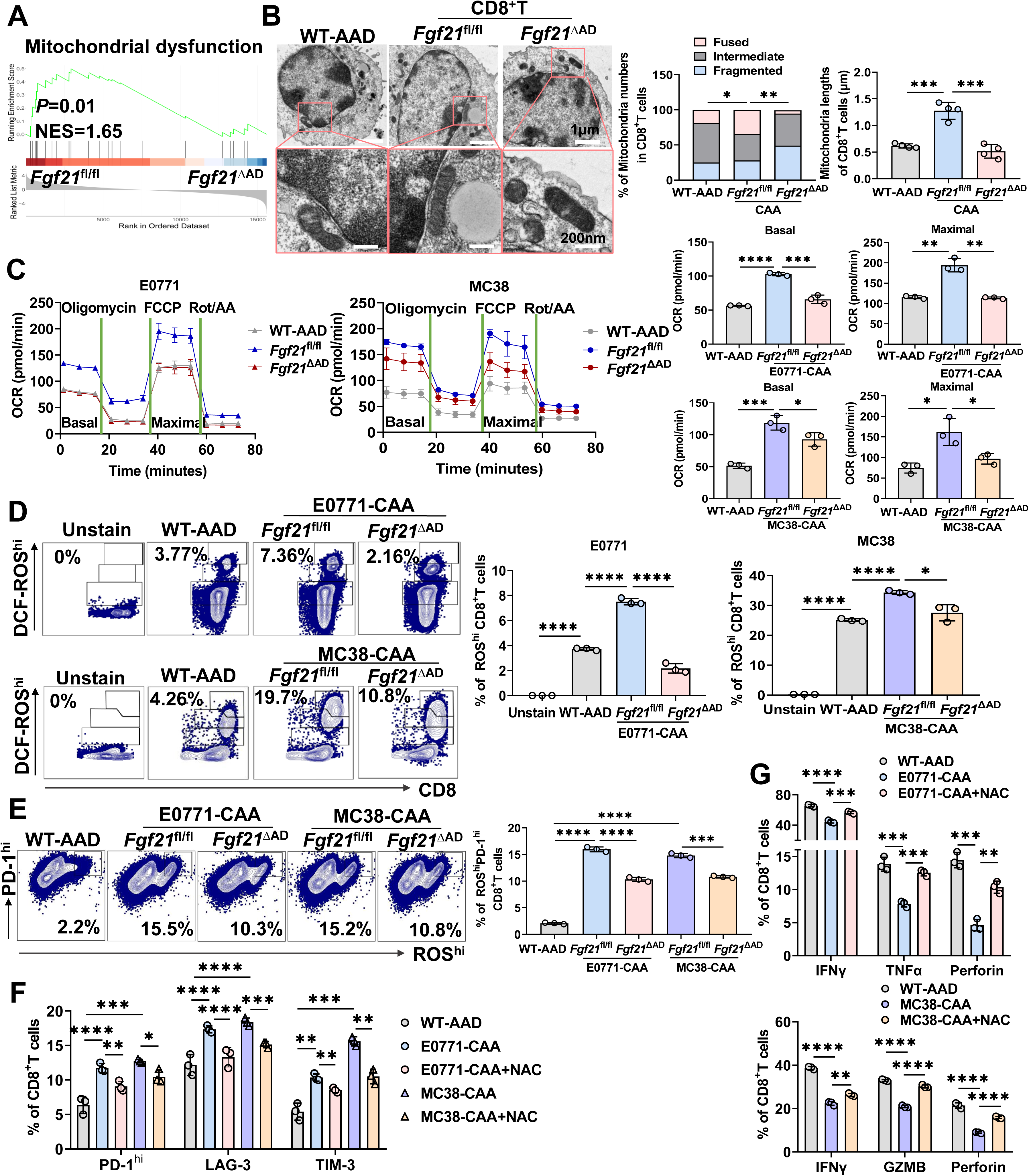
CAAs disturb mitochondrial homeostasis in exhausted CD8^+^T cells via FGF21. **A.** GSEA analysis of gene signatures of mitochondrial dysfunction of CD8^+^TILs sorted from MC38 tumors in *Fgf21*^fl/fl^ and *Fgf21*^ΔAD^ mice (n = 3 mice). **B.** Transmission Electron Microscope (TEM) analysis of mitochondrial morphology in CD8^+^T cells treated with conditioned media from WT-AAD or E0771 tumor-adjacent CAAs from *Fgf21*^fl/fl^ and *Fgf21*^ΔAD^ mice after activation by anti-CD3/CD28 for 96 hr (n = 4 biological replicates). **C.** Seahorse measurement of oxygen consumption rate (OCR) in CD8^+^T cells treated with conditioned media from WT-AAD or E0771/MC38 tumor-adjacent CAAs from *Fgf21*^fl/fl^ and *Fgf21*^ΔAD^ mice after activation by anti-CD3/CD28 for 96 hr (n = 3 biological replicates). **D.** FACS analysis of ROS^hi^CD8^+^T cells treated with conditioned media from WT-AAD or E0771/MC38 tumor-adjacent CAAs from *Fgf21*^fl/fl^ and *Fgf21*^ΔAD^ mice after activation by anti-CD3/CD28 for 96 hr (n = 3 biological replicates). **E.** FACS analysis of the percent of PD-1^hi^ROS^hi^ of CD8^+^T cells treated with conditioned media from WT-AAD or E0771/MC38 tumor-adjacent CAAs from *Fgf21*^fl/fl^ and *Fgf21*^ΔAD^ mice after activation by anti-CD3/CD28 for 96 hr (n = 3 biological replicates). **F.** FACS analysis of the percent of PD-1^hi^, LAG-3^+^, TIM-3^+^ of CD8^+^T cells treated with conditioned media from WT-AAD or E0771/MC38 tumor-adjacent CAAs, or plus with N-Acetylcysteine (NAC), after activation by anti-CD3/CD28 for 96 hr (n = 3 biological replicates). **G.** FACS analysis of the percent of IFNγ^+^, Granzyme B^+^, TNFα^+^ and Perforin^+^ of CD8^+^T cells treated with conditioned media from WT-AAD or E0771/MC38 tumor-adjacent CAAs, or plus with NAC, after activation by anti-CD3/CD28 for 96 hr (n = 3 biological replicates). Data are presented as the mean ± SEM. \**P*<0.05, \*\**P*<0.01, \*\*\**P*<0.001, \*\*\*\**P*<0.0001 by Student’s t test (B, C, D, E) and two-way ANOVA (F, G).

Considering the role of CAAs in promoting ROS production and inducing T cell exhaustion, we evaluated whether ROS scavenging with N-acetylcysteine (NAC) could reverse this effect. As expected, NAC treatment effectively abrogated CD8^+^T cell exhaustion and restored the cytotoxic molecule expression in CD8^+^T cells exposed to CAA-conditioned media (Figure 5F-G and S5D-F).

### FGF21 drives CAA lipolysis through ATGL via FGFR1-p38 signaling

We next sought to understand how FGF21 triggers CAA lipolysis. As mentioned above, lipolysis involves the hydrolysis of triglycerides by specific lipases, resulting in the liberation of FFAs. We speculated that FGF21 might modulate the expression of lipases to facilitate lipolysis. Indeed, GSEA analysis showed that genes linked to lipid lipolysis were significantly downregulated in the adipose tissue adjacent to MC38 tumors from *Fgf21*^ΔAD^ mice when compared to *Fgf21*^fl/fl^ mice (Figure 6A). Real-time PCR analysis further confirmed that FGF21 deficiency in adipose tissue significantly reversed the transcription of several lipolytic enzymes including ATGL, MGL, LAL, PNPLA3 (Figure 6B). ATGL is a key lipolytic enzyme that initiates the hydrolysis of triglycerides to release FFAs.^34,35^ Importantly, the protein expression of ATGL was dramatically increased in tumor-adjacent adipose tissue in contrast to normal adipose tissue, while FGF21 deletion remarkedly abrogated ATGL expression (Figure 6C), indicating that FGF21 enhances ATGL expression in CAAs. FGF21 activates the mitogen-activated protein kinase (MAPK) signaling pathway by binding to the FGFR1-KLB receptor complex.^36^ Western blot analysis revealed that p-FGFR1/KLB-p38 signaling was activated in CAAs. Yet, the absence of FGF21 evidently attenuated this signal axis (Figure 6D). Moreover, blockade of p-FGFR1/p38 signaling by using selective chemical inhibitors of FGFR1 and p38 notably suppressed ATGL expression (Figure 6E), suggesting that FGF21 upregulates ATGL expression through the FGFR1-p38 signaling pathway.

**Figure 6.**
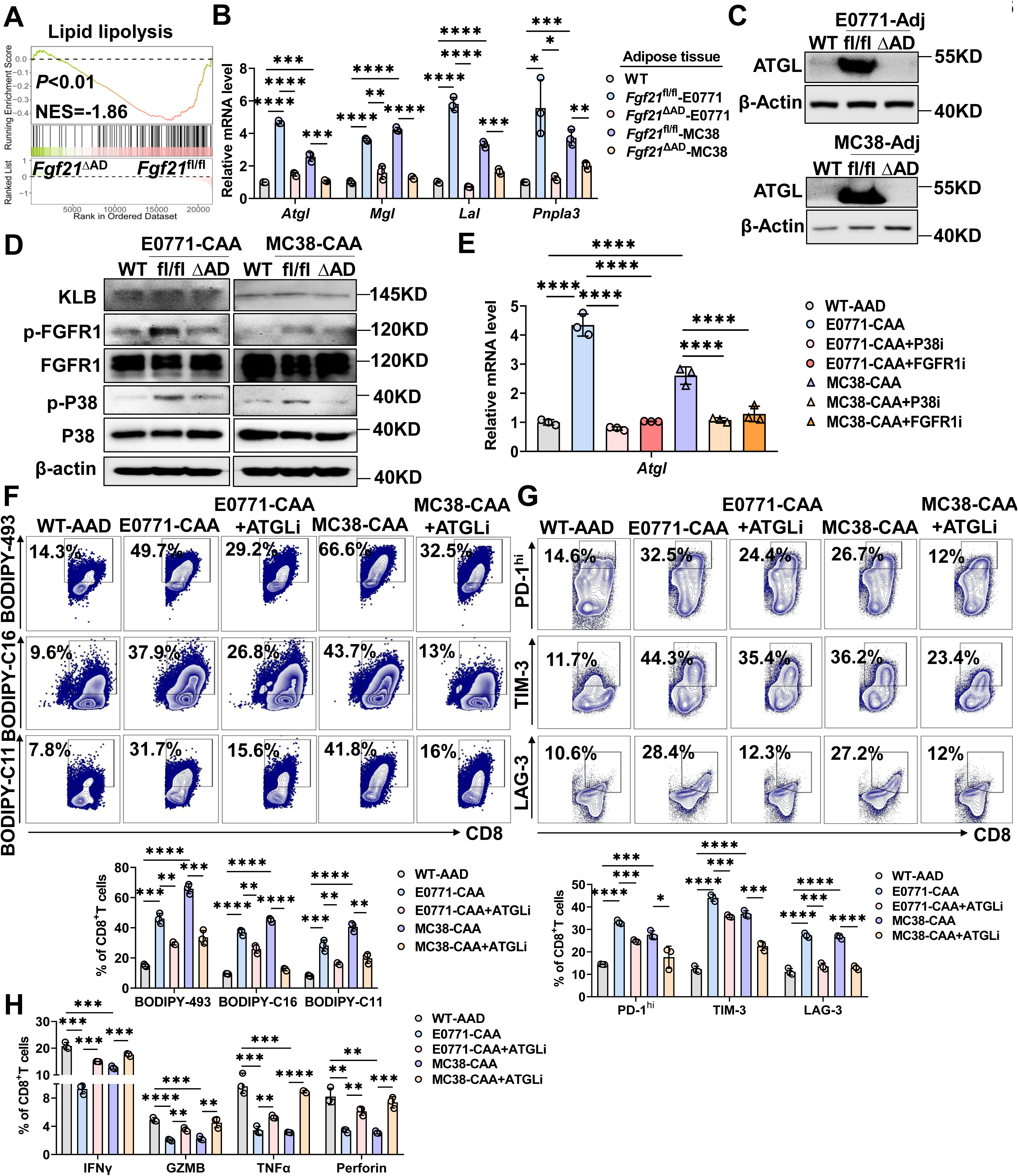
FGF21 drives CAA lipolysis through ATGL via FGFR1-p38 signaling. **A.** GSEA analysis of gene signatures of lipid lipolysis in MC38 tumor-adjacent adipose tissues from *Fgf21*^fl/fl^ and *Fgf21*^ΔAD^ mice (n = 3 mice). **B.** qPCR analysis of gene expression of lipolytic lipases in WT normal adipose tissues and E0771/MC38 tumor-adjacent adipose tissues from *Fgf21*^fl/fl^ and *Fgf21*^ΔAD^ mice (n = 3 mice). **C.** Western Blot analysis of ATGL expression in WT normal adipose tissues and E0771 tumor-adjacent adipose tissues from *Fgf21*^fl/fl^ and *Fgf21*^ΔAD^ mice (n = 3 independent experiments). **D.** Western Blot analysis of p-FGFR1/FGFR1, KLB, p-p38/p38 in WT-AAD, and *Fgf21*^fl/fl^ and *Fgf21*^ΔAD^ transwell CAAs educated by E0771/MC38 tumor cells for 6 hr (n = 3 independent experiments). **E.** qPCR analysis of *Atgl* expression in WT-AAD, E0771/MC38 tumor cell-educated CAAs, plus with P38 inhibitor and FGFR1 inhibitor treatment. **F.** FACS analysis of BODIPY-493, BODIPY-C16 and BODIPY-C11 staining in CD8^+^T cells treated with conditioned media from WT-AAD or E0771/MC38 tumor cell-educated CAAs, plus with ATGL inhibitor treatment, after activation by anti-CD3/CD28 for 96 hr (n = 3 biological replicates). **G.** FACS analysis of the percent of PD-1^hi^, LAG-3^+^ and TIM-3^+^ of CD8^+^T cells treated with conditioned media from WT-AAD or E0771/MC38 tumor cell-educated CAAs, plus with ATGL inhibitor treatment, after activation by anti-CD3/CD28 for 96 hr (n = 3 biological replicates). **H.** FACS analysis of the percent of IFNγ^+^, Granzyme B^+^, TNFα^+^ and Perforin^+^ of CD8^+^ T cells treated with conditioned media from WT-AAD or E0771/MC38 tumor cell-educated CAAs, plus with ATGL inhibitor treatment, after activation by anti-CD3/CD28 for 96 hr (n = 3 biological replicates). Data are presented as the mean ± SEM. \**P*<0.05, \*\**P*<0.01, \*\*\**P*<0.001, \*\*\*\**P*<0.0001 by Student’s t test (E) and two-way ANOVA (B, F, G, H).

To further clarify the role of ATGL-driven CAA lipolysis in CD8^+^T cell lipid uptake and function, we employed the ATGL inhibitor Atglistatin to disrupt CAA lipolysis. Of note, ATGL inhibition markedly reduced lipid uptake and peroxidation in CD8^+^T cells treated with CAA-conditioned media (Figure 6F). Consequently, the expression of inhibitory receptors on CD8^+^T cells decreased (Figure 6G), while the expression of cytotoxic molecule was greatly enhanced (Figure 6H and Figure S6), indicating that suppression of lipolysis in CAAs by ATGL inhibitor recovered CD8^+^T cell effector function.

Taken together, all these data suggest that ATGL is a critical downstream target of FGF21-FGFR1-p38 signal axis to drive CAA lipolysis, consequently mediating CD8^+^T cell dysfunction.

### Combinational treatment with ATGL inhibitor and anti-PD-1 synergistically improves the anti-tumor immune response

Since targeting CAA lipolysis with Atglistatin improves CD8^+^T cell function in *vitro*, we aimed to explore whether Atglistatin administration in *vivo* could enhance the therapeutic efficacy of ICIs. Notably, the combination of Atglistatin and anti-PD-1 therapy resulted in a substantial delay in tumor growth (Figure 7A) and significant tumor size reduction compared with monotherapy groups (Figure S7A). We further evaluated the immune responses in tumors post treatments. We observed a significant increase in the infiltration of CD8^+^T cells in tumors within the combination treatment group (Figure 7B and S7B). Furthermore, the combination treatment markedly reduced lipid uptake and peroxidation in CD8^+^TILs (Figure 7C and Figure S7C), coupled with a notable decrease in T cell exhaustion (Figure 7D and S7D) and more potent production of cytotoxic molecules (Figure 7E and S7E). These data demonstrate that combination of ATGL inhibitor and PD-1 blockade potentiates CD8^+^T cell anti-tumor immune response and synergistically blunts tumor growth.

**Figure 7.**
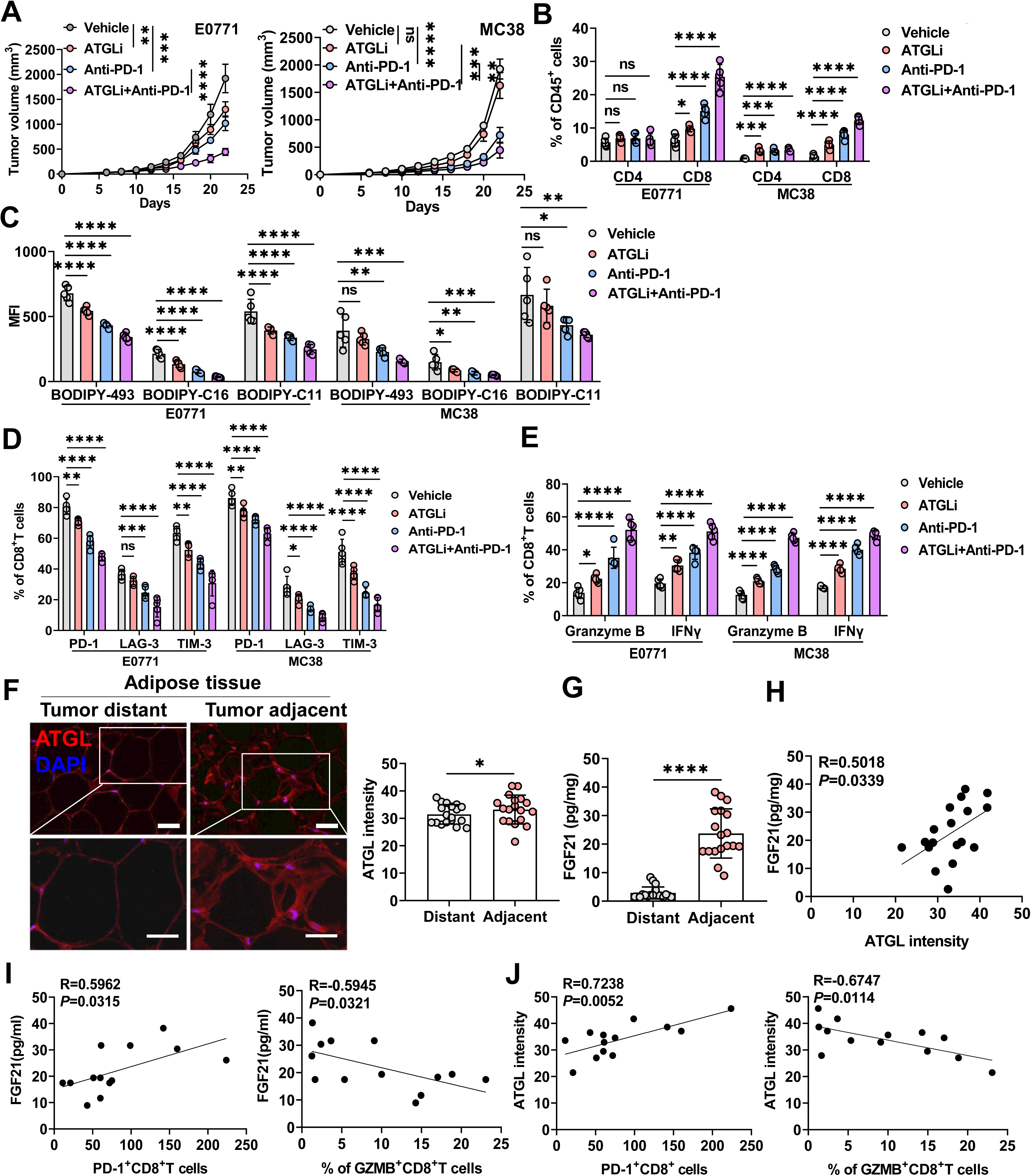
Combinational treatment with ATGL inhibitor and anti-PD-1 synergistically improves the anti-tumor immune response. **A.** Tumor growth curve of E0771 and MC38 tumors treated with Vehicle, ATGL inhibitor, anti-PD-1, or ATGL inhibitor combined with anti-PD-1 (n = 5 mice). **B.** FACS analysis of the percent of tumor-infiltrated CD4^+^, CD8^+^T cells in E0771 and MC38 tumors after treatment with Vehicle, ATGL inhibitor, anti-PD-1, or ATGL inhibitor combined with anti-PD-1 (n = 5 mice). **C.** FACS analysis of BODIPY-493, BODIPY-C16 and BODIPY-C11 staining of CD8^+^TILs in E0771 and MC38 tumors after treatment with Vehicle, ATGL inhibitor, anti-PD-1, or ATGL inhibitor combined with anti-PD-1 (n = 5 mice). **D.** FACS analysis of PD-1, LAG-3 and TIM-3 in CD8^+^TILs in E0771 and MC38 tumors after treatment with Vehicle, ATGL inhibitor, anti-PD-1, or ATGL inhibitor combined with anti-PD-1 (n = 5 mice). **E.** FACS analysis of IFNγ and Granzyme B in CD8^+^TILs in E0771 and MC38 tumors with Vehicle, anti-ATGL, anti-PD-1, or ATGL inhibitor combined with anti-PD-1 (n = 5 mice). **F.** IF staining of ATGL in tumor-adjacent and distant adipose tissues from breast cancer patients (n=18 patients). **G.** ELISA analysis of FGF21 protein levels in tumor-adjacent and distant adipose tissues from breast cancer patients (n=18 patients). **H.** Linear regression analysis of the correlation of FGF21 and ATGL expression in tumor-adjacent adipose tissues from breast cancer patients (n=18 patients). **I.** Linear regression analysis of the correlation of FGF21 levels in tumor-adjacent adipose tissues and the number of CD8^+^PD-1^+^ cells or CD8^+^Granzyme B^+^ cells in tumor tissues (n=13 patients). **J.** Linear regression analysis of the correlation of ATGL expression in tumor-adjacent adipose tissue and the number of CD8^+^PD-1^+^ cells or CD8^+^Granzyme B^+^ cells in tumor tissues (n=13 patients). Data are presented as the mean ± SEM. \**P*<0.05, \*\**P*<0.01, \*\*\**P*<0.001, \*\*\*\**P*<0.0001 by Student’s t test (F, G), two-way ANOVA (A, B, C, D, E) and liner regression analysis (H, I, J).

Finally, we determined the clinical relevance of FGF21-ATGL-driven CAA lipolysis and CD8^+^TIL dysfunction in breast cancer patients. We observed that tumor-adjacent adipose tissue exhibited higher levels of FGF21 and ATGL expression compared to the normal distant adipose tissue (Figure 7F-G). Moreover, FGF21 expression was positively correlated with ATGL expression in the tumor-adjacent adipose tissue (Figure 7H). Importantly, both FGF21 and ATGL expression correlated positively with the number of PD-1^+^CD8^+^T cells, but negatively with Granzyme B^+^CD8^+^T cells (Figure 7I-J and S7F). These results suggest that increased FGF21 and ATGL expression in tumor-adjacent adipose tissues contribute to CD8^+^TIL exhaustion and impaired functionality in breast cancer patients.

## Discussion

Tumor cells reprogram various neighboring cells, including fibroblasts, immune cells, adipocytes and other stromal cells, to support their own proliferation, invasion and metastasis.^37^ For a long time, adipocytes are often ignored for their influence within TME. However, their role in fueling cancer is becoming increasingly evident in recent years.^38,39^ A groundbreaking study by Dirat and colleagues reveals the crucial role of peritumoral adipose tissue in breast cancer progression. They first demonstrate that tumor cells can reprogram nearby adipocytes to develop unique characteristics.^5^ Tumor cell-adipocyte coculture reduces the expression of adipocyte differentiation markers, coinciding with increased expression of proinflammatory cytokines. Persistent contact between adipocytes and tumor cells make them to acquire a fibroblast-like morphology.^40,41^ In many solid tumors, the invasion of tumor cells into the proximal adipose tissue leads to profound delipidation and activation of adipocytes. These cancer-associated adipocytes (CAAs) exchange cytokines and lipids with tumor cells, leading to their metabolic reprogramming, consequently promoting tumor cell proliferation and invasion.^42–44^ Here we validate that CAAs lose the morphology of normal mature adipocytes, lowering lipid storage. Through transcriptomics and metabolomics analysis, we further demonstrate that CAAs undergo lipolysis and release FFAs, indicating that CAAs contribute to the lipid enrichment within TME.

Despite the interactions between CAAs and tumor cells have been well recognized, TME also involves the infiltration of immune cells, particularly CD8^+^T cells, which are key effector cells in anti-tumor immune responses.^45^ Bioinformatic analyses have shown that in patients with triple-negative breast cancer, high adipocyte generation is associated with low CD8^+^T cell infiltration and immune suppression.^46^ Similarly, in triple-negative breast cancer tumor models, CXCL8 derived from CAAs inhibits CD8^+^T cell infiltration and upregulates PD-L1 expression on tumor cells.^47^ Besides, in obesity-related breast cancer models, leptin secreted by adipocytes induces CD8^+^T cell fatty acid oxidation via activation of STAT3 signaling pathway, thereby suppressing CD8^+^T cell effector function.^48^ These previous studies indicate that CAAs mediate immune suppression. However, the mechanism underlying CAA lipolysis and how CAAs regulate CD8^+^T cell lipid metabolism remain unclear. Here we identify FGF21 is a key factor driving CAA lipolysis and FFA releasing, which promotes the lipid uptake and peroxidation in CD8^+^T cells. The lipid accumulation in CD8^+^T cells disturbs mitochondrial homeostasis and causes ROS overproduction, eventually resulting in T cell exhaustion. Moreover, we reveal that FGF21 triggers CAA lipolysis by upregulating ATGL, which is a critical lipase initiating lipid hydrolysis. Genetic deletion of FGF21 in adipocytes or ATGL inhibition dampens CAA lipolysis, leading to reduced lipid uptake by CD8^+^T cells and restoring their anti-tumor immunity. Our findings uncover the critical role of FGF21-ATGL-mediated lipolysis in CAAs in limiting CD8^+^T cell anti-tumor responses, suggesting that targeting CAA lipolysis could be a promising strategy to enhance the cancer immunotherapy efficacy.

FGF21 is an endocrine hormone that senses metabolic stress and balances energy homeostasis. FGF21 can be locally released from adipose tissue, where it acts as an autocrine factor in adipocytes to increase glucose uptake and promote thermogenesis.^26,49^ FGF21 regulates lipolysis in adipocytes in response to the metabolic state. Under normal feeding, FGF21 stimulates lipolysis in white adipose tissue, but inhibits it during fasting.^50^ Besides, acute FGF21 treatment has been found to suppress lipolysis in primary adipocytes.^51^ Here we report that CAAs secret higher levels of FGF21 compared to WT adipocytes. FGF21 autocrinally promotes lipolysis in CAAs, resulting in the release of FFAs, which in turn reprogram the lipid metabolism of CD8^+^TILs. The release of FFAs from adipocyte lipolysis appears to further increase FGF21 expression,^52^ suggesting a potential feedback loop in which FGF21 facilitates CAA lipolysis and FFA release. Additionally, our previous research has shown that tumor cells abnormally secret FGF21.^28^ We cannot exclude the possibility that tumor-secreted FGF21 may also contribute to CAA lipolysis. ATGL is the key enzyme for the release of fatty acids from triacylglycerol stores during intracellular lipolysis.^53^ Indeed, our results indicate a significant upregulation of ATGL in tumor-adjacent adipose tissue compared to normal adipose tissue. Moreover, FGF21 expression is positively correlated with ATGL expression in tumor-adjacent adipose tissue of breast cancer patients. FGF21 exerts its effects by stimulating FGFR1/2 in conjunction with the coreceptor KLB, and has been shown to activate ΑΚΤ-mTORC1 and ERK signaling via MAPK in adipocytes.^24,54^ Our results reveal that FGF21 upregulates ATGL expression through FGFR1/KLB-p38 signaling activation in CAAs. In line with this, in nonalcoholic fatty liver disease, liver-derived FGF21 has been reported to induce hepatic p38 activation, increasing the influx of fatty acids from adipose tissue to liver, which contributes to hepatic ectopic lipid accumulation.^55^ Further studies are warranted to delineate the precise molecular mechanism by which FGF21 regulates ATGL expression. Importantly, disrupting CAA lipolysis by the ATGL inhibitor Atglistatin robustly enhances CD8^+^T cell anti-tumor immune response, thereby improving the therapeutic efficacy of anti-PD-1 therapy. Atglistatin treatment has been report to prevent myocardial damage,^56^ improve cardiac function,^57^ and alleviate lipotoxicity and inflammation.^58,59^ Our findings highlight the potential of Atglistatin as a therapeutic agent to augment cancer immunotherapy.

CD8^+^T cells are critical for the elimination of malignant cells. Within TME, both intrinsic and extrinsic metabolic cues contribute to CD8^+^T cell dysfunction.^60^ One key immunosuppressive feature of TME is lipid accumulation. Imbalance in lipid uptake or synthesis affect CD8^+^T cell effector function.^16,61^ For instance, intrapancreatic CD8^+^T cells accumulate specific long-chain fatty acids, which dampens their anti-tumor function by decreasing mitochondrial activity.^19^ CD36-mediated uptake of fatty acids by CD8^+^TILs induces lipid peroxidation and ferroptosis, leading to reduced cytotoxic cytokine production and impaired anti-tumor activity.^20,21^ Fatty acid utilization in mitochondria induces oxidative stress and impacts mitochondrial dynamics, which refers to the process of fission, fusion, mitophagy and transport.^62^ Mitochondrial fission supports quality control by removing damaged mitochondria, while fusion facilitates the exchange of intramitochondrial components, helping to maintain mitochondrial integrity.^63^ An imbalance in mitochondrial dynamics can compromise mitochondrial function, leading to abnormal cellular fate.^64^ Loss of the mitochondrial fission protein dynamic-related protein 1 (DRP1) has been shown to negatively affect TIL function by accelerating exhaustion.^65^ PD-1 signaling inhibits T cell activation through reducing DRP-dependent mitochondrial fission. In MC38 tumors, CD8^+^T cells expressing PD-1 exhibit reduced DRP activities and elongated mitochondria compared to PD-1 negative counterparts.^66^ Our results reveal that CAA-enhanced fatty acid oxidation in CD8^+^T cells impairs their mitochondria dynamics, as evidenced by increased mitochondrial fusion and elongation in cells exposed to CAA-conditioned media. This fusion exacerbates OXPHOS and induces excessive ROS, driving CD8^+^T cell exhaustion. However, CD8^+^T cells treated with conditioned media from FGF21-deficient CAAs show a reversal of mitochondrial fusion and normalization of mitochondrial dynamics, coinciding with restored cytotoxic function. These findings support that mitochondrial dynamics are essential for optimizing T cell anti-tumor activity.

### Limitations of the Study

This study has several limitations. First, it is unclear whether FGF21-driven CAA lipolysis is influenced by the body’s metabolic state, such as starvation, cold exposure or obesity. Future studies are warranted to examine whether these parameters affect FGF21 regulation of CAA lipolysis. Secondly, this study primarily focuses on the interaction between CAAs and CD8^+^T cells. Since the TME contains various immune cell types, we cannot rule out the possibility that fatty acids released by CAAs may affect the function of other immune cells, thereby indirectly impact the anti-tumor activity of CD8^+^T cells. Lastly, we have observed the upregulation of several lipolytic enzymes, including ATGL, HSL and MGL, in tumor-adjacent adipose tissue. While we mainly explore the mechanism by which FGF21 induces ATGL upregulation, the mechanism underlying the upregulation of other lipolytic enzymes remain unclear. Further investigations in this area will be helpful for refining strategies to target CAA lipolysis and enhance the therapeutic efficacy of cancer immunotherapy.

## Supporting information

Graphical Abstract

## Acknowledgements

This work was supported by funds from National Natural Science Foundation of China NSFC 82472915 (J.G.), 32422029 (J.G.), 82103518 (J.G.), 82272997 (H.L.), Innovative research team of high-level local Universities in Shanghai SHSMU-ZDCX20210802 (J.G.), Shanghai Jiao Tong University School of Medicine "Clinical Full-time Research Team" project (J.G.). “NSFC Promotion Project” of Renji Hospital RJTJ23-RC-005 (J.G.). State Key Laboratory of Systems Medicine for Cancer Program KF2414 (S.D.).

## Author contributions

J.G. conceived the study and designed the research. S.D., C.H., W.Q., H.L., R.Z., L.L., P.L. performed the experiments. C.Y., W.Y., J.L. collected clinical samples. X.L., X.Y. did bioinformatic analysis. J.G., S.D., C.H. interpreted data. J.G., S.D. wrote the manuscript. All the authors discussed and read the manuscript.

## Materials and Methods

### Human samples

The human samples were collected from 18 breast cancer patients who received total mastectomy. The tumor-adjacent adipose tissues (<2 cm to the tumor site) and distant adipose tissues (the farthest distance to the tumor site) were collected from the species. These adipose tissue samples were snap-freezing in liquid nitrogen for RNA and Protein analysis, or frozen in Tissue-Tek O.C.T. compound for cryosection. For H&E staining, samples were fixed in 4% Paraformaldehyde (PFA) solution for paraffin embedding. Tumor samples were collected from 13 patients from this cohort and fixed in 4% PFA for immunofluorescence staining of CD8, PD-1 and Granzyme B. The clinical characteristics of all these patients were described in **Table S1**. The informed content was obtained from all study participants. The study was approved by the Ethics Committee of the Renji Hospital, Shanghai Jiao Tong University School of Medicine.

### Mice

Mice in six-to eight-week old were used in our experiments and housed under specific pathogen free (SPF) conditions in the animal facility at Ren Ji Hospital. WT C57BL/6 mice were purchased from Shanghai Model Organisms Center. *Adipoq*-Cre (C001042, C57BL/6N) and *Fgf21*^flox/+^ mice (CKOCMP-56636-Fgf21-B6N-VA, C57BL/6N) were purchased from Cyagen Biosciences. Inc, and crossed to generate *Fgf21*^fl/fl^ mice and *Adipoq*-Cre;*Fgf21*^fl/fl^ mice (*Fgf21*^ΔAD^). All these mice were viable and fertile with no reported abnormalities. The genotyping PCR primers were provided in **Table S2.** Littermate animals from different cages were randomly assigned into experimental groups. All the animal studies were conducted in accordance with the protocols approved by the Animal Care and Use Committee at Renji Hospital, Shanghai Jiao Tong University School of Medicine.

### Cell lines

E0771 mammary adenocarcinoma cell line was purchased from ATCC. MC38 murine colorectal cell line was kindly given by Dr. Bin Ma (School of Biomedical Engineering and Med-X Research Institute, Shanghai Jiao Tong University). E0771 were cultured in RPMI-1640 (Gibco) and MC38 were cultured in DMEM (Gibco). All cells were maintained in the media supplemented with 10% fetal calf serum (FBS, Gibco) and 100 U/ml penicillin/streptomycin (Gibco) at 37°C under 5% CO_2_ incubator. All of the cell lines were regularly tested for *Mycoplasma* free.

### Animal tumor models

For syngeneic transplanted breast tumor model, 2*10^6^ E0771 tumor cells suspended in 100 μl PBS were orthotopically injected into the fat pad of 4^th^ mammary gland of WT, *Fgf21*^fl/fl^, or *Fgf21*^ΔAD^ female mice. For colorectal tumor model, 1*10^6^ MC38 tumor cells suspended in 100 μl PBS were subcutaneously (s.c.) injected into the right flank of WT, *Fgf21*^fl/fl^, or *Fgf21*^ΔAD^ mice. Tumor volumes were measured along largest diameter and perpendicular diameter (a, b) and calculated as V=a*b*b/2. When the tumor volume reached about 1000 mm^3^, mice were euthanized, and the tumor tissues and tumor-adjacent adipose tissues were collected.

For in *vivo* treatments, ATGL inhibitor Atglistatin was dissolved in 10% dimethyl sulfoxide (DMSO), 40% PEG 300, 5% Tween 80 and 45% PBS. Mice with palpable tumors were intraperitoneally administrated with 400 μmol/kg Atglistatin every day (cat#S7364, Selleck). Anti-PD-1 mAb (cat#BE0146, Bio X Cell) or the isotype control (BE0089#, Bio X Cell) was intraperitoneally administrated with 100 μg/mouse every four days.

### Primary adipocytes differentiation

The primary adipocyte isolation and differentiation in *vitro* was conducted based on the protocol from Paul Cohen’s lab.^67^ Briefly, adipose tissues were mechanically minced and enzymatically digested for 35 min at 37°C with buffer containing 2 mg/ml Collagenase IV (cat#5138, Sigma), 2.4 mg/ml Dispase II (cat#40104ΕS80, Yeasen) and 20 mM CaCl_2_ in a 50 ml falcon tube. The digested solution was filtered by a 70 μm cell strainer, then spin down at 1500 rpm for 10 min. The red blood cells were lysed by Lysing Buffer (cat#555899, BD) for 3 min. Cells were resuspended with ΒenchStable^ΤΜ^ DMEM/F12 medium containing 10% FBS, and plated into 12-well gelatin (cat#40108ΕS60, Yeasen) coated plates. Pre-adipocytes were differentiated after the cells reached 100% confluence with adipocyte differentiation cocktail mixture containing 0.5 mM IBMX (cat#Ι5879, Sigma), 1 μM Dexamethasone (cat#D4902, Sigma), 850 nM Insulin (cat#Ι8830, Solarbio); 1 μM Rosiglitazone (cat#R2408, Sigma). On day 2, media was switched to 850 nM insulin and 1 μM rosiglitazone. On day 4, rosiglitazone was removed, media containing insulin (850 nM) was replaced every other day. On day 6, adipocytes were fully differentiated. The conditioned media (CM) of adipocytes were harvest 24 hr after culturing with FBS-free DMEM/F12 media. Pre-adipocytes isolated from subcutaneous adipose tissue of naïve WT mice were differentiated and used as control.

### Tumor cells educate adipocytes by Transwell

Preadipocytes isolated from WT tumor-free mice were prepared as described above. After six-day cocktail culture, the cells successfully differentiated into mature adipocytes. Subsequently, 5*10^5^ E0771 or MC38 tumor cells were seeded into 12-mm transwell chamber (cat#3460, 0.4 μm pore, Costar) to educate the mature adipocytes. For the analysis of adipocyte phenotype and lipid content, a 48 hr education period was conducted. For Western blot analysis of signaling activation by tumor cells, a 6 hr education period was performed. For the blockade of FGF21-activated signaling in adipocytes, 100 nM FGFR1 inhibitor PD173074 (cat#S1264, Selleck), 1 μM p38 inhibitor SB203580 (cat#S1076, Selleck), or 100 nM ATGL inhibitor Atglistatin (cat#S7364, Selleck) was added during education.

### Oil Red O staining

The *in vitro* differentiated adipocytes were fixed with 4% PFA at 4°C for 30 min, then washed with PBS three times. For the staining of human adipose tissues, O.C.T embedded tissues were cryosectioned into 30 μm section. The slide sections were fixed with 4% PFA at 4°C for 1 hr, then washed with PBS for three times. Cells or slides were stained with 3 g/L Oil Red O (cat#O0625, Sigma) at 37°C for 1 hr. After washing with PBS for three times, images were captured by a fluorescence microscope (OLYMPUS) under white light.

### BODIPY-493 immunofluorescence staining

The slide sections of human adipose tissue were fixed with 4% PFA at 4°C for 1 hr, then washed with PBS for three times. After that, the slides were permeabilized by buffer (cat#88-8823-88, eBioscience) at 4°C for 1 hr. After washing with PBS for three times, the slides were stained with 2 μM BODIPY-493/508 (cat#D3922, Invitrogen) at 37°C dark incubator for 30 min. The stained slides were imaged by a fluorescence microscope (OLYMPUS).

### LC-MS analysis

The supernatant collected from WT AADs or MC38 CAAs were subjected to global metabolomics profiling using an ultrahigh-performance liquid chromatography–mass spectrometry (UPLC–MS) platform following conventional metabolomics procedures by Sangon Βiotech (Shanghai) Co., Ltd. Significant lipid alterations between the WT AAD and CAA group were identified based on Variation Importance in the Projection (VIP) values (>1) and absolute Log_2_ fold change (FC) (>1). VIP values were extracted from the OPLS-DA results generated using the R package MetaboAnalystR. To avoid overfitting, a permutation test (200 permutations) was performed. Identified lipids were annotated using the KEGG compound database.

### Free fatty acid quantification assay

The free fatty acid levels in the supernatant of WT-AAD and CAAs were measured by by Free Fatty Acid Quantitation Kit (cat#MAK044, Sigma). Briefly, 0, 2, 4, 6, 8, and 10 μl of the standard solution was added into a 96 well plate. Fatty Acid Assay Buffer was added into each well to bring the volume to 50 μl. Samples were added with Fatty Acid Assay Buffer to the final volume of 50 μl. 2 μl ACS (Acyl-CoA Synthetase) reagent was added into each sample and the standard wells, then incubated for 30 minutes at 37 °C. After that, 50 μl Master Reaction Mix (Fatty Acid Assay Buffer 44 μl, Fatty Acid Probe 2 μl, Enzyme Mix 2 μl, Enhancer 2 μl) was added into each well and incubated for 30 min at 37 °C in dark. For colorimetric assay, the absorbance was measured at 570 nm (A570). The concentration of Fatty Acids was calculated by Fa/Sv = C (Fa = amounts of fatty acids in unknown sample (nmole) from standard curve; Sv = sample volume (μl) added to reaction well; C = concentration of fatty acids in sample).

### Enzyme-linked Immunosorbent Assay (ELISA)

The protein expression of FGF21 in the supernatant of WT AADs and CAAs, or the adipose tissue lysates from patients were measured by mouse (Sangon Biotech, cat# D721010-0096) or human ELISA kit (Sangon Biotech, cat# D711036-0096) as previously described.^28^ The protein level was calculated based on the standard curve.

### Quantitative reverse transcription PCR

Total RNA from adipocytes or adipose tissues was extracted using Trizol (Invitrogen) according to the manufacturer’s instructions and reversed transcribed with HiScript III All-in one RT SuperΜix Perfect for qPCR kit (cat#R333-01, Vazyme) to make cDNA. Real-time PCR was performed using SYBR qPCR Master Mix (cat#Q711-03, Vazyme). The expression of each gene was calculated based on the cycle threshold which set within the linear range of DNA amplification. The relative expression was calculated by the cycle threshold method, with normalization of raw data to a housekeeping gene gapdh (2-^ΔΔ^Ct). The primer sequences are provided in **Table S2**.

### Western Blotting

Adipose tissues were grinded into powder in liquid nitrogen, then lysed with RIPA buffer containing protease inhibitor cocktail. Mouse adipocytes educated by E0771/MC38 tumor cells for 6 hr were lysed with RIPA buffer. The whole-cell lysates were boiled for 10 min and loaded on SDS-PAGE gels, electrophoresed, transferred onto PVDF membranes, blocked with 5% BSA-TBST or nonfat milk-TBST, and further incubated with the primary and secondary antibodies. After that, the membranes were washed with TBST for 5 times. The blots were developed using ECL reagent with Bio-Rad ChemiDoc Touch machine. The following antibodies were used: anti-FGFR1 (cat#9740, CST, 1:1000); PhosphoFGFR1 (cat#52928, CST, 1:1000); anti-KLB (cat#AF2619, R&D, 1:1000); p38 MAPK (D13E1) (cat#8690, CST, 1:1000); Phospho-p38 MAPK (Thr180/ Tyr182) (cat#4511, CST, 1:1000); β-Actin (cat#S0B0074, STARTER, 1:1000); ATGL (cat#2138, CST, 1:1000); HRP-Rabbit anti-Goat (cat#AS029, Abclonal, 1:20000); HRP Goat anti-Rabbit IgG (H+L) (cat#AS014, Abclonal,1:20000).

### CD8^+^T cell isolation and treatment with conditioned media

Naïve CD8^+^ T cells were isolated from mouse splenocytes using EasySep Mouse CD8^+^T Cell Isolation Kit (cat#19858, STEMCELL). Naive CD8^+^T cells were seeded into 96-well round plate which was precoated with anti-CD3 antibody (10 μg/ml), and stimulated with anti-CD28 antibody (2 μg/ml), containing IL-2 (2 ng/ml). T cells were treated with conditioned media (CM) from adipocytes plus complete media (1:1) during activation. Cells were harvested at different times points and conduced flow cytometry analysis.

### Flow cytometry analysis

For the staining of tumor-infiltrating immune cells, tumors were immediately dissected after mice were sacrificed. Tumors were finely minced and incubated in DMEM (Gibco) supplemented with collagenase IV (cat#C5138, Sigma) at 1.5 mg/mL, and DNase I (cat#11284932001, Roche) at 100 mg/mL on a shaker at 250 rpm, 37°C for 45 min. Complete medium with FBS was added to the mixture to terminate digestion, and the mixture was filtered with a 70 μm cell strainer, washed with PBS once, and resuspended with 1% BSA-PBS containing 1 mM EDTA. The cells were incubated with anti-mouse CD16/32 antibody for 15 min on ice to block nonspecific Fc receptor binding. Cells were then stained with cell surface markers: anti-CD45-APC/Cyanine7 (cat#103116, Clone 30-F11, Biolegend), anti-CD11c-Bv510 (cat#117337, Clone N418 Biolegend), anti-I-A/I-E-Bv650 (cat#107641, Clone M5/114/15/2, Biolegend), anti-CD103-PE/Cyanine7 (cat#121425, Clone 2E7, Biolegend), anti-CD11b-APC (cat#101212, M1/70, Biolegend), anti-CD11b-Bv605 (cat#3101237, Clone M1/70, Biolegend), anti-Ly-6C-PE (cat#128008, Clone HK1.4, Biolegend), anti-Ly-6G-APC/Cyanine7 (cat#127624, Clone 1A8, Biolegend), anti-F4/80-PerCP/Cyanine5.5 (cat#123126, Clone BM8, Biolegend), anti-CD3-APC (cat#100236, Clone 17A2, Biolegend), anti-CD8-PE (cat#126608, Clone YTS156.7.7, Biolegend), anti-CD4-PerCP/Cyanine5.5 (cat#100434, Clone GK1.5, Biolegend) and incubated on ice for 30 min. Live cells were stained by eFluor 455 (cat#65-0868, eBioscience). After washing with PBS, cells were resuspended with FACS buffer and acquired by LSR Fortessa flow cytometry (BD Biosciences). Data were analyzed with Flowjo software (V10, BD). For intracellular staining of T cell cytotoxic molecules, cells were pre-incubated with Cell Stimulation Cocktail (cat#00-4970-93, eBioscience) and Brefeldin A (cat#88-8823-88, eBioscience) for 6 hr. After staining with surface markers: anti-CD45-APC/Cyanine7 (cat#103116, Clone 30-F11, Biolegend), anti-CD3-AF700 (cat#100216, Clone17A2, Biolegend), anti-CD8-PE (cat#126608, CloneYTS156.7.7, Biolegend), cells were washed with PBS and fixed with fixation buffer (cat#88-8823-88, eBioscience) for 30 min. After that, cells were stained with anti-IFNγ-APC (cat#554413, CloneXMG1.2, BD), anti-Granzyme B-Percpcy5.5 (cat#372212, CloneQA16A02, Biolegend), anti-Perforin-Pacific Blue (cat#154311, CloneS16009A, Biolegend), anti-TNFα-PE (cat#506305, CloneMP6-XT22, Biolegend) antibodies diluted in permeabilization buffer for 30 min. Cells were washed with PBS and acquired by LSR Fortessa flow cytometry.

For the intracellular staining of lipid contents in T cells, cells were washed with PBS and fixed with fixation and permeabilization buffer (cat#88-8823-88, eBioscience) for 30 min. After that, cells were stained with 2 μM BODIPY-493/508 (cat#D3922, Invitrogen), 2 μM BODIPY-FL-C16 (cat#D3821, Invitrogen) or 2 μM BODIPY-581/591-C11 (cat#D3861, Invitrogen) for 30 min at 37°C in the dark. Cells were washed and resuspended with FACS buffer and acquired by LSR Fortessa flow cytometry.

For T cell exhaustion maker staining, the following antibodies were used: anti-PD-1-APC/Cy7 (cat#135224, Clone29F.1A12, Biolegend), anti-TIM-3-PE/Cy7 (cat#134009, CloneB8.2C12, Biolegend), anti-LAG3-Percp/Cyanine5.5 (cat#125211, CloneC9B7W, Biolegend).

### Seahorse assay in CD8^+^T cells

CD8^+^T cells were exposed to the conditioned media from WT AADs and CAAs during anti-CD3/CD28 activation. Seahorse XF Cell Mito Stress Test Kit (cat#103010-100, Agilent) was used to measure the oxygen consumption rate (OCR) on a XFe96 Analyzer (Agilent Seahorse). 2*10^5^ CD8^+^T cells were seeded in a XFe96 microplates (cat#103793-100, Agilent). After treatment, cells were washed with XF assay media for one time and then incubated in the media for 1 hr. Before detection, cells were spin down for adhering at 300 g for 2 min. After measuring the basal OCR, 2.5 μM oligomycin, 2.5 μM FCCP, 0.2 μM Rot/AA were added in real time, and the maximal OCR was measured.

### ROS detection

CD8^+^T cells were cultured with conditioned media from WT AADs and CAAs during anti-CD3/CD28 activation. Cells were harvested at different time points, and stained with anti-CD3-APC, anti-CD8-PE, DCFH-DA probe (cat#S0033S, Beyotime). After washing with PBS, cells were resuspended with FACS buffer and acquired by LSR Fortessa flow cytometry (BD Biosciences). Data were analyzed with Flowjo software. For ROS scavenging, 2 mM N-acetylcysteine (cat#HYB0215, MCE) was added when the CD8^+^T cells exposed to the conditioned media from CAAs.

### Transmission Electron Microscopy

CD8^+^T cells were cultured with conditioned media from WT AADs, *Fgf21*^fl/fl^ E0771-CAAs or *Fgf21*^ΔΑD^ E0771-CAAs for 96 hr. Cells were fixed with electron microscope fixative buffer (cat#G1102, Servicebio), post-fixed in 1% osmic acid solution, dehydrated with a graded ethanol series, pure acetone and embedded in embedding agent. Sectioning was performed with LEICA EM Journal Pre-proof UC7 ultrathin microtome (Leica, EM UC7) and imaging were performed by Scientific Compass platform.

### Η&Ε and immunofluorescence staining

The tumor tissues and adipose tissues harvested from breast cancer patients were fixed in 4% PFA overnight, embedded in paraffin, and sectioned for H&E and immunofluorescent staining (IF). For IF staining, the sections underwent deparaffinization in xylene, sequential rehydration through ethanol, and PBS rinsing. Antigen retrieval was carried out by microwaving for 30 min in a specialized antigen retrieval solution, followed by natural cooling to room temperature. To block non-specific signals, 5% goat serum in PBS was applied. Slides were incubated with anti-ATGL (cat#2138, CST, 1:200), anti-CD8 (cat#ab178089, Abcam, 1:200), anti-PD-1 (cat#86163, CST, 1:100), or anti-Granzyme B (ab255598#, Abcam, 1:200) overnight at 4°C. After that, slides were washed and further incubated with the secondary antibody Goat anti-Rabbit IgG (H+L) Alexa Fluor™488 (cat#A-11008, Invitrogen, 1:1000), or Goat anti-Rabbit IgG (H+L) Alexa Fluor™647 (cat#A-21244, Invitrogen, 1:1000) for 1 hr at room temperature. Finally, the coverslips were mounted on microscope slides with Prolong Antifade with DAPI. Fluorescent images were captured using the Olympus A1Si Laser Scanning Confocal Microscope. The cell area of adipose tissue and IF intensity was analyzed by Image J.

### RNA-sequencing

Normal adipose tissues were collected from WT tumor-free mice. Tumor-adjacent adipose tissues were collected from MC38 tumor-bearing mice (*Fgf21*^fl/fl^ and *Fgf21*^ΔΑD^), and snap freezing in liquid nitrogen. For CD8^+^TIL RNA sequencing, the single-cell suspension of tumor tissues from MC38 tumor-bearing mice (*Fgf21*^fl/fl^ and *Fgf21*^ΔΑD^) was prepared as described above. After staining with anti-CD45, anti-CD3, and anti-CD8 antibodies, CD8^+^T cells were sorted out by BD FACS Aria (BD Biosciences). Both the adipose tissues and sorted cells were extracted RNA with Trizol. RNA Sequencing were performed by Sangon Βiotech Co., Ltd (Shanghai). Paired-end read sequences were aligned to the mouse reference genome (version mm10) using the default settings in STAR (version 2.6.1b) and quantified by HTSeq (version 0.11.0) in ‘‘intersection-strict’’ mode. Differentially expressed genes were (DEG) identified with log-fold change>1.5 and false discovery rate (FDR)<0.05. The expression differences from mouse transcriptome data were assessed by logarithm base two-fold changes of expression (log_2_FC). The gene functional enrichment was performed by gene set enrichment analysis (GSEA). The enrichment magnitude and statistical significance were quantified by normalized enrichment score (NES) and *P* value. The RNA-sequencing data has been deposited to Sequence Read Archive (SRA) data under accession code PRJNA1170421.

### Quantification and Statistical Analysis

All described results are representative of at least three independent experiments. The number of samples (n) were described in detail for each figure panel. No statistical method was used to predetermine sample size. The experiments were not randomized, except that the mice were randomly grouped before treatment. Investigators were not blinded to allocation during the experiments and outcome assessments. Data were presented as individual values. Statistical analysis was performed using GraphPad prism 10.0 software. Two-tailed unpaired Student’s t-test was used for the comparison between two groups. One-way analysis of variance (ANOVA) or two-way ANOVA followed by the Sidak’s test was used for the multiple comparisons. Repeated-measures two-way ANOVA (mixed model) followed by the Sidak’s multiple comparisons test was used for analysis of the tumor growth curve. Statistical significance was denoted with *(*P* < 0.05), **(*P* < 0.01), ***(*P* < 0.001), **** (*P*<0.0001) in the figures.

---

**Figure S1.**
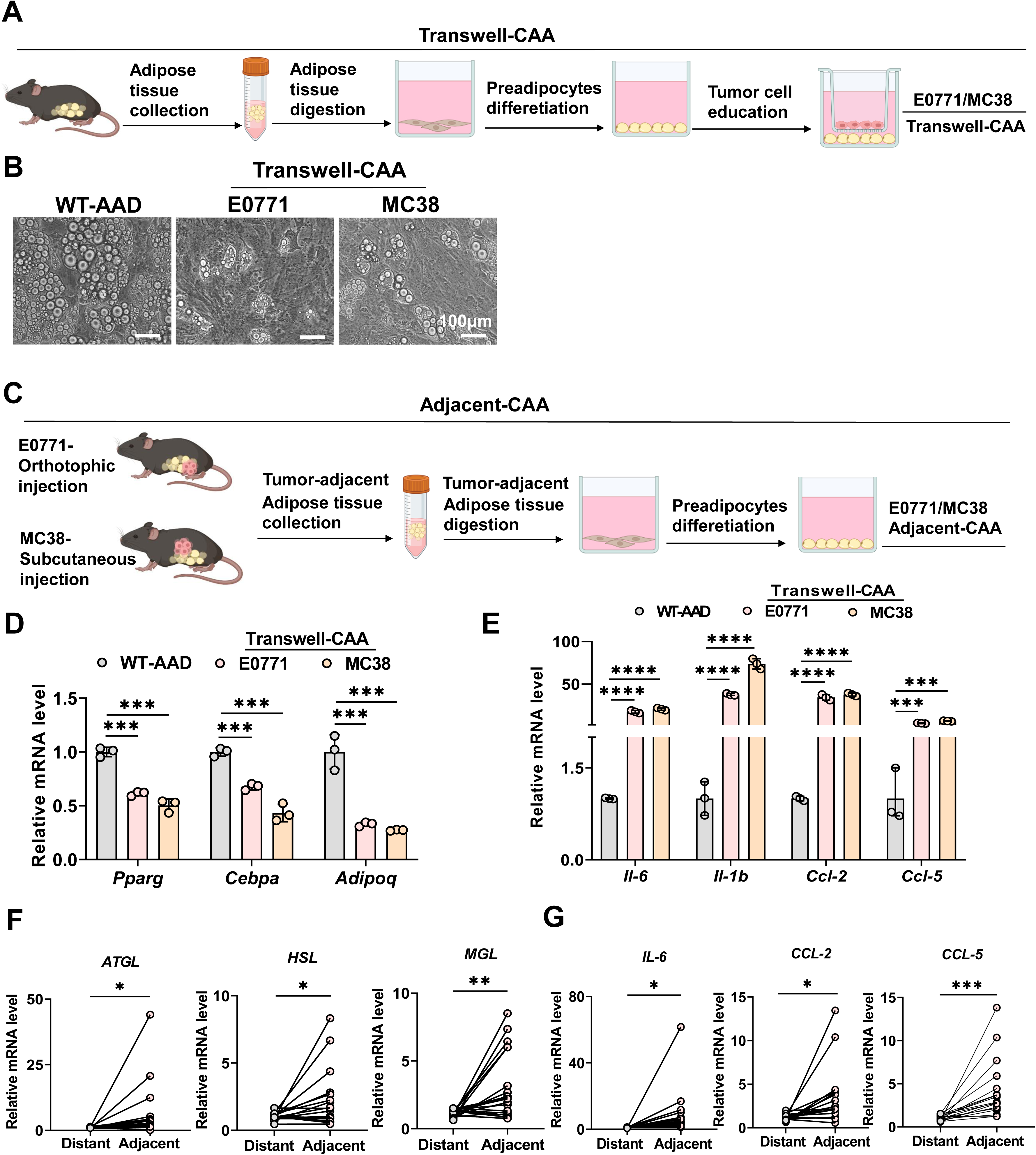
Cancer-associated adipocytes (CAAs) exhibit delipidation, related to Figure 1. **A.** Schematic diagram for tumor cell-educated adipocytes by transwell (Transwell-CAA). **B.** Adipocyte morphology of WT AADs and E0771/MC38 Transwell CAAs. Scale bar: 100 μm. **C.** Schematic diagram for tumor-adjacent adipocyte generation (Adjacent-CAA). **D.** qPCR analysis of gene expression of mature adipocyte markers, including PPARγ, CEBPa, Adiponectin, in WT AADs and E0771/MC38 Transwell CAAs (n = 3 biological replicates). **E.** qPCR analysis of proinflammatory gene expression, including IL-6, IL-b, CCL-2, CCL-5, in WT AADs and E0771/MC38 Transwell CAAs (n = 3 biological replicates). **F.** qPCR analysis of gene expression of lipolytic lipases, including ATGL, HSL, MGL, in the tumor-adjacent and distant adipose tissues from breast cancer patients (n = 18 patients). **G.** qPCR analysis of proinflammatory gene expression, including IL-6, CCL-2, CCL-5 in the tumor-adjacent and distant adipose tissues from breast cancer patients (n = 18 patients). Data are presented as the mean±SEM. \**P*<0.05, \*\**P*<0.01, \*\*\**P*<0.001, \*\*\*\**P*<0.0001 by Student’s t test (F, G) and two-way ANOVA (D, E).

**Figure S2.**
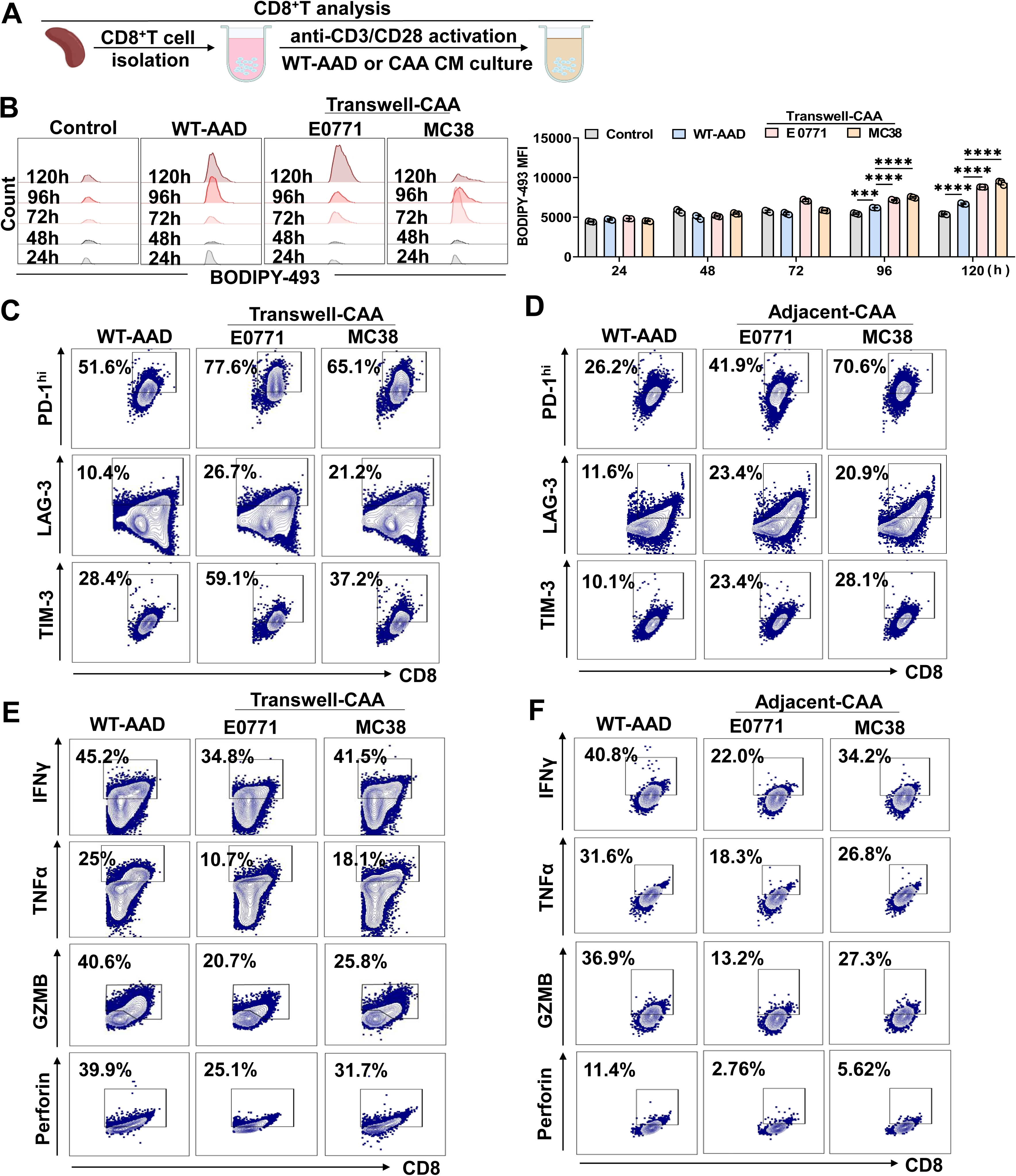
CAAs mediate CD8^+^T cell dysfunction, related to Figure 2. **A.** Schematic diagram for CD8^+^T cells treated with conditioned media (CM) from WT-AADs, Transwell-CAAs or adjacent-CAAs during anti-CD3/CD28 activation. **B.** FACS analysis of BODIPY-493 staining in CD8^+^T cells treated with conditioned media from WT-AADs or Transwell-AADs at different time points during anti-CD3/CD28 activation (n = 3 biological replicates). **C-D.** Representative FACS analysis of PD-1, LAG-3, and TIM-3 staining in CD8^+^T cells treated with conditioned media from WT-AADs, Transwell-CAAs **(C)** or adjacent-CAAs **(D)**. **E-F.** Representative FACS analysis of IFNγ, TNFα, Granzyme B, and Perforin staining in CD8^+^T cells treated with conditioned media from WT-AADs, Transwell-CAAs **(E)** or adjacent-CAAs **(F)**. Data are presented as the mean±SEM. \*\*\**P*<0.001, \*\*\*\**P*<0.0001 by two-way ANOVA (B).

**Figure S3.**
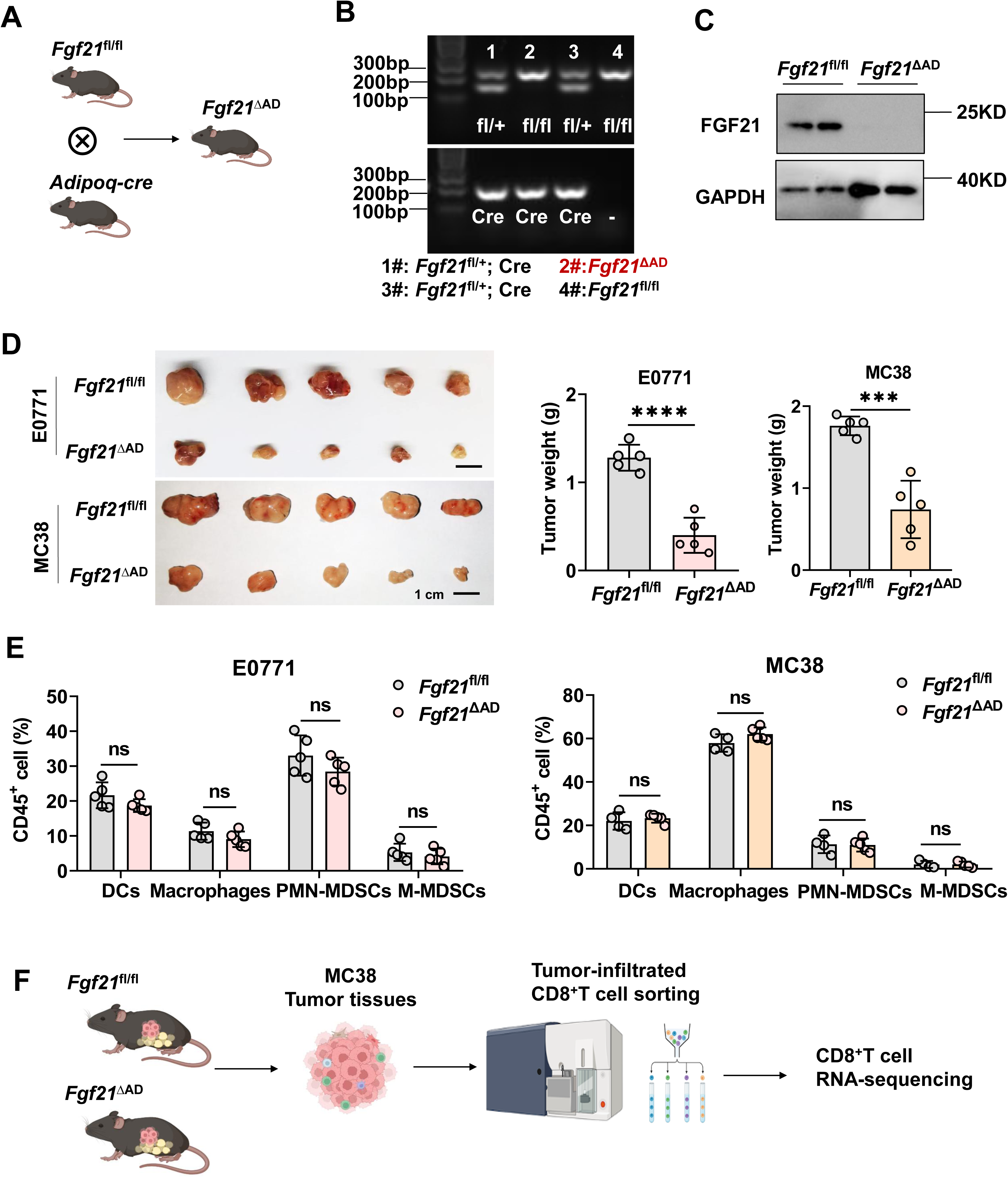
Generation of adipose tissue specific *Fgf21* knock out mice, related to Figure 3. **A.** Schematic diagram for *Fgf21*^fl/fl^ mice crossing with *Adipoq*-cre mice to generate adipose tissue specific *Fgf21* knock out mice (*Fgf21*^ΔAD^). **B.** Genotyping results for *Fgf21*^fl/fl^, *Fgf21*^fl/+^, *Fgf21*^fl*/+*^*; Adipoq*-Cre, *Fgf21*^fl/fl^*; Adipoq*-Cre (*Fgf21*^ΔAD^) mice. **C.** Western blot analysis of FGF21 protein expression in the adipose tissues of *Fgf21*^fl/fl^ and *Fgf21*^ΔAD^ mice (n = 3 independent experiments). **D.** Images of tumor mass and tumor weight of E0771 and MC38 in *Fgf21*^fl/fl^ and *Fgf21*^ΔAD^ mice (n = 5 mice). **E.** FACS analysis of the percent of DCs, macrophages, PMN-MDSCs and M-MDSCs in E0771 and MC38 tumors of *Fgf21*^fl/fl^ and *Fgf21*^ΔAD^ mice (n = 4-5 mice). **F.** Schematic diagram for CD8^+^TIL RNA-sequencing. Data are presented as the mean±SEM. \*\*\**P*<0.001, \*\*\*\**P*<0.0001, ns, no significance by Student’s t test (D) and two-way ANOVA (E).

**Figure S4.**
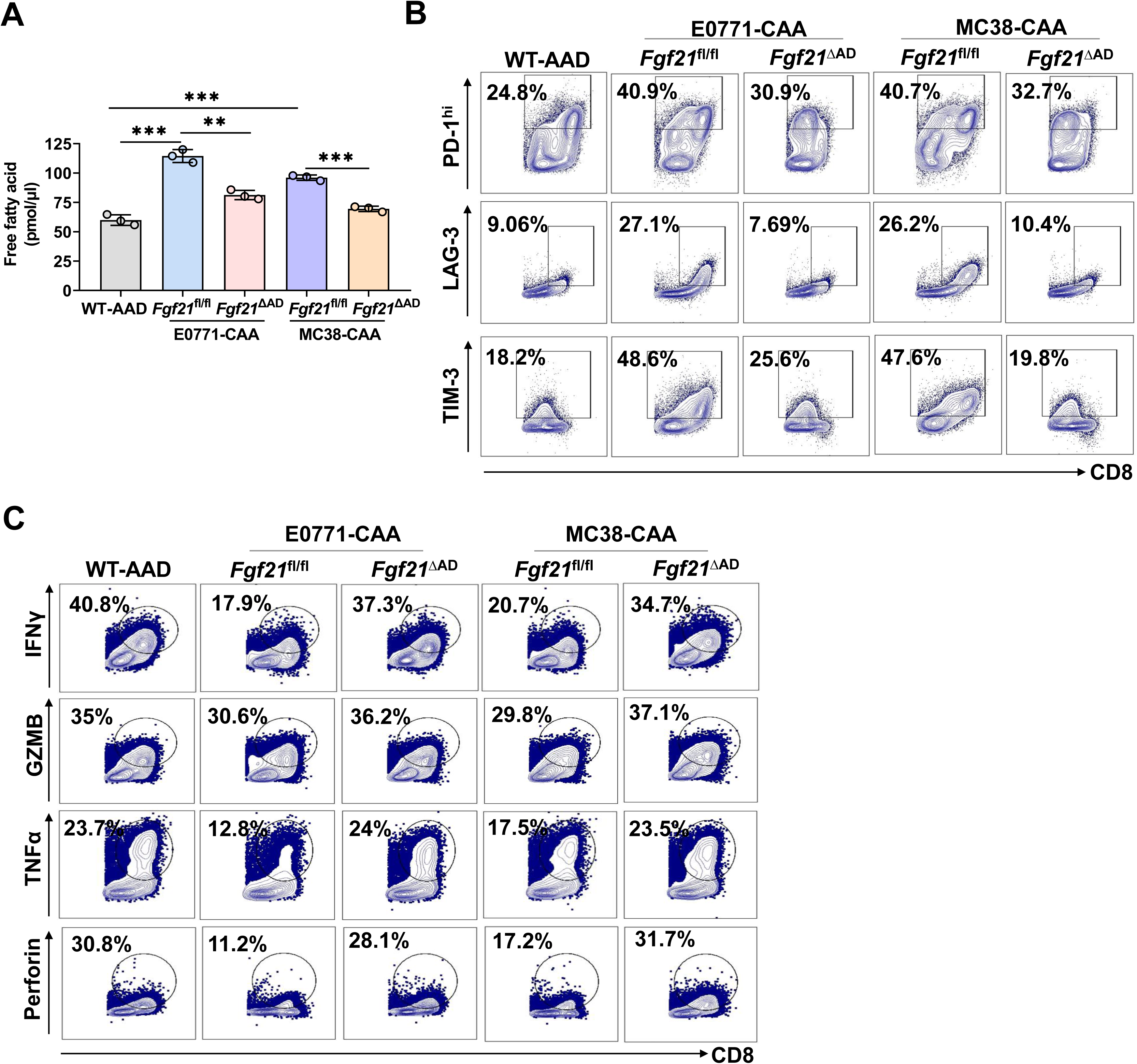
FGF21 deficiency in adipocytes restores cytotoxic function of CD8^+^T cells exposed to CAA conditioned media, related to Figure 4. **A.** Free fatty acid levels in the supernatant of WT-AADs and E0771/MC38 tumor-adjacent CAAs from *Fgf21*^fl/fl^ and *Fgf21*^ΔAD^ mice (n = 3 biological replicates). **B.** Representative FACS analysis of PD-1, LAG-3 and TIM-3 staining in CD8^+^T cells treated with conditioned media from WT-AADs or E0771/MC38 tumor-adjacent CAAs from *Fgf21*^fl/fl^ and *Fgf21*^ΔAD^ mice after activation by anti-CD3/CD28 for 96 hr. **C.** Representative FACS analysis of IFNγ, Granzyme B, TNFα and Perforin staining in CD8^+^T cells treated with conditioned media from WT-AADs or E0771/MC38 tumor-adjacent CAAs from *Fgf21*^fl/fl^ and *Fgf21*^ΔAD^ mice after activation by anti-CD3/CD28 for 96 hr. Data are presented as the mean±SEM. \**P*<0.05, \*\**P*< 0.01, \*\*\**P*<0.001 by Student’s t test (A).

**Figure S5.**
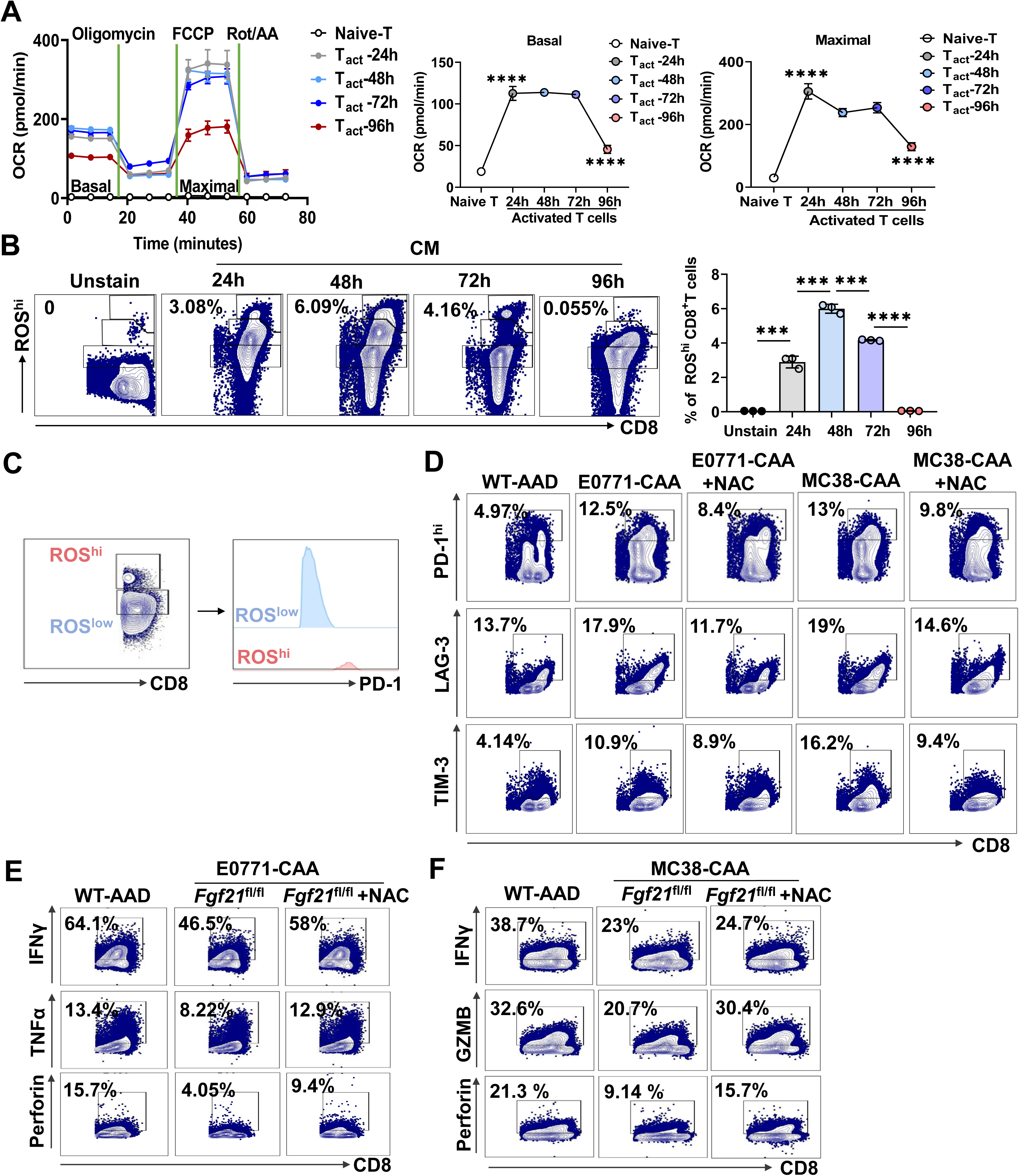
CAAs aggravate ROS production in CD8^+^T cells, related to Figure 5. **A.** Seahorse measurement of oxygen consumption rate (OCR) in CD8^+^T cells at different time points during activation by anti-CD3/CD28 (n = 3 biological replicates). **B.** FACS analysis of ROS^hi^CD8^+^T cells at different time points during activation by anti-CD3/CD28 (n = 3 biological replicates). **C.** FACS analysis of PD-1 expression of ROS^hi^ and ROS^low^CD8^+^T cells. **D.** Representative FACS analysis of PD-1, LAG-3, TIM-3 staining in CD8^+^T cells treated with conditioned media from WT-AADs or E0771/MC38 tumor-adjacent CAAs, or plus with N-Acetylcysteine (NAC), after activation by anti-CD3/CD28 for 96 hr. **E-F.** Representative FACS analysis of IFNγ, Granzyme B, TNFα and Perforin staining in CD8^+^T cells treated with conditioned media from WT-AADs or E0771/MC38 tumor-adjacent CAAs, or plus with N-Acetylcysteine (NAC), after activation by anti-CD3/CD28 for 96 hr. Data are presented as the mean±SEM. \*\*\**P*<0.001, \*\*\*\**P*<0.0001, ns, no significance by Student’s t test. (A, B)

**Figure S6.**
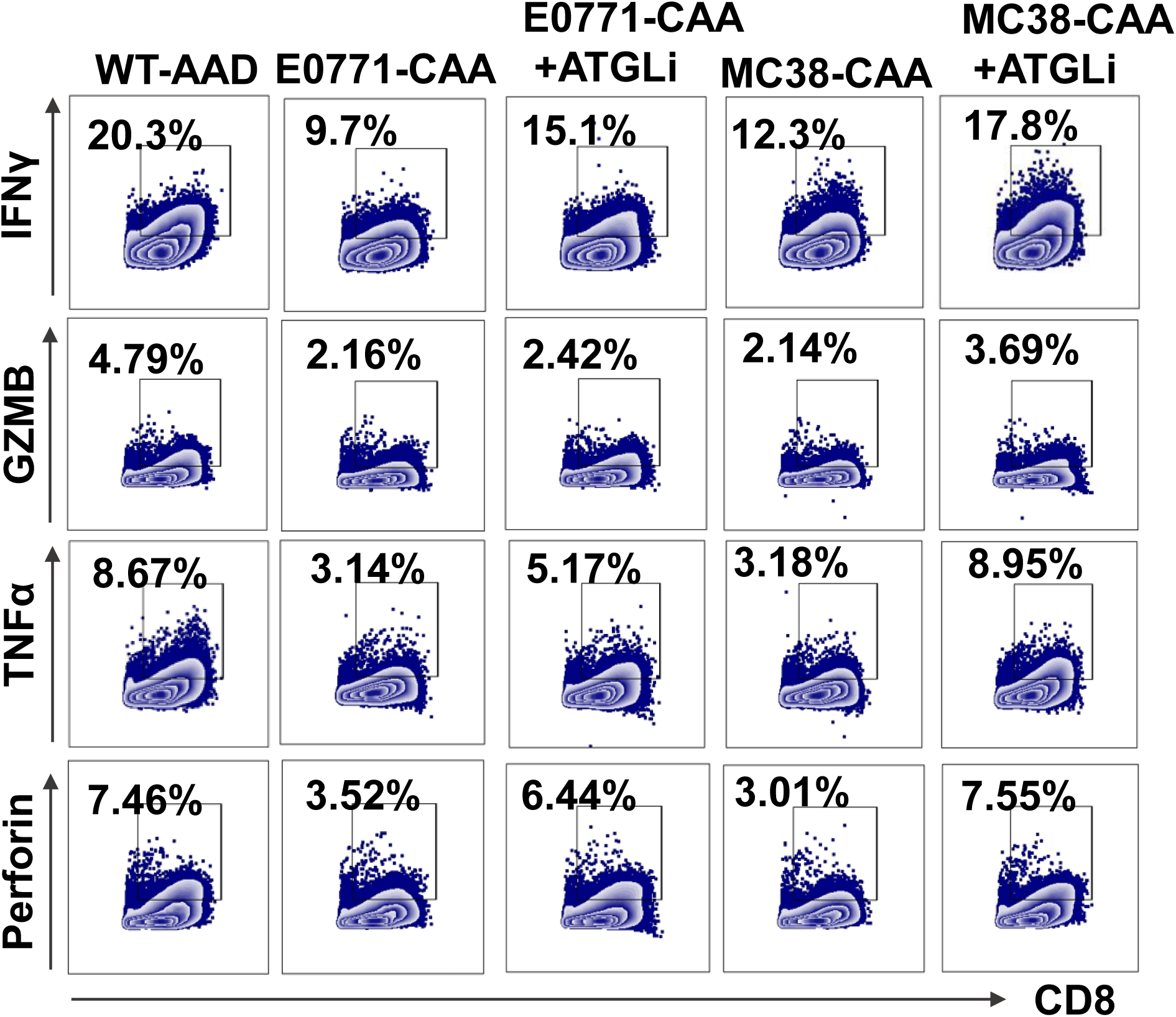
ATGL inhibitor treatment restores CD8^+^T cell function in *vitro*, related to Figure 6. Representative FACS analysis of IFNγ, Granzyme B, TNFα and Perforin staining in CD8^+^T cells treated with conditioned media from WT-AADs or E0771/MC38 tumor cell-educated CAAs, plus with ATGL inhibitor treatment, after activation by anti-CD3/CD28 for 96 hr.

**Figure S7.**
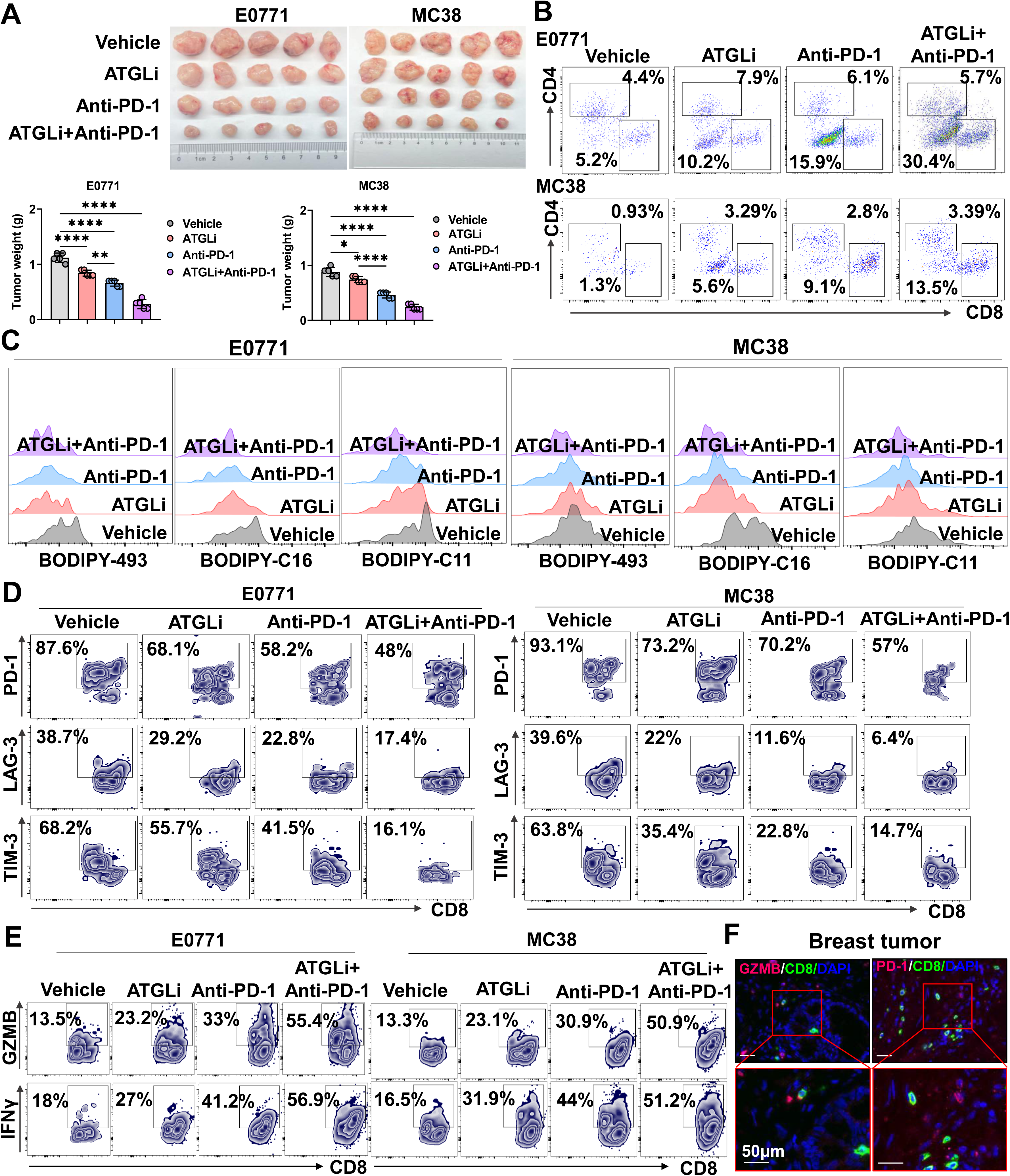
Combinational treatment of ATGL inhibitor and anti-PD-1 enhances the anti-tumor activity of CD8^+^T cells, related to Figure 7. **A.** Images of tumor mass and tumor weight of E0771 and MC38 in the mice with Vehicle, ATGL inhibitor, anti-PD-1, or ATGL inhibitor combined with anti-PD-1 treatment (n = 5 mice). **B.** Representative FACS analysis of tumor-infiltrated CD4^+^ and CD8^+^T cells in E0771 and MC38 tumors after treatment with Vehicle, ATGL inhibitor, anti-PD-1, or ATGL inhibitor combined with anti-PD-1. **C.** Representative FACS analysis of BODIPY-493, BODIPY-C16 and BODIPY-C11 staining in CD8^+^TILs of E0771 and MC38 tumors after treatment with Vehicle, ATGL inhibitor, anti-PD-1, or ATGL inhibitor combined with anti-PD-1. **D.** Representative FACS analysis of PD-1, LAG-3 and TIM-3 staining in CD8^+^TILs of E0771 and MC38 tumors after treatment with Vehicle, ATGL inhibitor, anti-PD-1, or ATGL inhibitor combined with anti-PD-1. **E.** Representative FACS analysis of IFNγ, Granzyme B staining of in CD8^+^TILs of E0771 and MC38 tumors after treatment with Vehicle, ATGL inhibitor, anti-PD-1, or ATGL inhibitor combined with anti-PD-1. **F.** Representative IF staining of CD8^+^, Granzyme B^+^ and PD-1^+^ cells in tumor tissues of breast cancer patients. Scale bar: 50 μm. Data are presented as the mean±SEM. \**P*<0.05, \*\**P*<0.01, \*\*\**P*<0.001, \*\*\*\**P*<0.0001 by Student’s t test (A).

