## Supplementary material for "Cancer-associated adipocytes mediate CD8^+^T cell dysfunction via FGF21-driven lipolysis": Table S1

| Patient | Sex | Age (years) | Menopause |
|---|---|---|---|
| 1 | Female | 67 | Yes |
| 2 | Female | 60 | Yes |
| 3 | Female | 65 | Yes |
| 4 | Female | 70 | Yes |
| 5 | Female | 66 | Yes |
| 6 | Female | 43 | No |
| 7 | Female | 53 | No |
| 8 | Female | 70 | Yes |
| 9 | Female | 56 | Yes |
| 10 | Female | 40 | No |
| 11 | Female | 45 | No |
| 12 | Female | 47 | No |
| 13 | Female | 60 | Yes |
| 14 | Female | 54 | No |
| 15 | Female | 48 | No |
| 16 | Female | 62 | Yes |
| 17 | Female | 70 | Yes |
| 18 | Female | 57 | Yes |

| Pathology | Tumor Size(cm) | Axillary LN |
|---|---|---|
| Invasive carcinoma | 1.5 | Negative |
| Apocrine differentiated carcinoma | 2 | Negative |
| Invasive carcinoma | 2 | Negative |
| Invasive carcinoma | 2 | Negative |
| Invasive carcinoma | 2.3 | Negative |
| Invasive carcinoma | 3 | 8/24 |
| Invasive carcinoma | 2 | Negative |
| Invasive carcinoma | 2.5 | 1/15 |
| Invasive carcinoma | 1 | Negative |
| Invasive carcinoma | 1.5 | Negative |
| Invasive carcinoma | 2 | Negative |
| Ductal carcinoma in situ | 1.5 | Negative |
| Invasive carcinoma | 2 | 6/13 |
| [illegible]uctal carcinoma with microinvasio[illegible] | 2.5 | Negative |
| Invasive carcinoma | 3 | Negative |
| Invasive carcinoma | 1.5 | Negative |
| Mucinous adenocarcinoma | 3.5 | Negative |
| Invasive carcinoma | 2 | 2/16 |

| N stage | Metastasis | M stage | TNM stage | Stage |
|---|---|---|---|---|
| 0 | No | 0 | T1N0 | IA |
| 0 | No | 0 | T1N0 | IA |
| 0 | No | 0 | T1N0 | IA |
| 0 | No | 0 | T1N0 | IA |
| 0 | No | 0 | T2N0 | IIA |
| 2 | No | 0 | T2N2 | IIIA |
| 0 | No | 0 | T1N0 | IA |
| 1 | No | 0 | T2N1 | IIB |
| 0 | No | 0 | T1N0 | IA |
| 0 | No | 0 | T1N0 | IA |
| 0 | No | 0 | T1N0 | IA |
| 0 | No | 0 | T1N0 | IA |
| 2 | No | 0 | T2N2 | IIIA |
| 0 | No | 0 | T2N0 | IIA |
| 0 | No | 0 | T2N0 | IIA |
| 0 | No | 0 | T1N0 | IA |
| 0 | No | 0 | T2N0 | IIA |
| 1 | No | 0 | T1N1 | IIA |

| ER(%) | PR(%) | HER2 Status | Ki67(%) |
|---|---|---|---|
| 95 | 1 | - | 10 |
| 0 | 0 | + | 30 |
| 90 | 80 | - | 30 |
| 90 | 90 | - | 30 |
| 0 | 0 | - | 40 |
| 30 | 40 | - | 10 |
| 90 | 90 | - | 30 |
| 70 | 0 | - | 15 |
| 90 | 0 | - | 10 |
| 90 | 90 | - | 15 |
| 0 | 0 | - | 80 |
| 0 | 0 | - | 10 |
| 95 | 95 | - | 10 |
| 0 | 0 | + | 40 |
| 70 | 80 | - | 20 |
| 90 | 90 | - | 10 |
| 95 | 90 | - | 10 |
| 0 | 0 | + | 60 |
