## Supplementary material for "Cancer-associated adipocytes mediate CD8^+^T cell dysfunction via FGF21-driven lipolysis": Table S2

**Table S2. Primer sequences**

| Primers for Real-time PCR | | |
|---|---|---|
| Gene ID | Species | Sequence (5' -> 3') |
| *Pparg* | Mouse | F' GGAAGACCACTCGCATTCCTT<br>R' GTAATCAGCAACCATTGGGTCA |
| *Cebpa* | Mouse | F' GCGGGAACGCAACAACATC<br>R' GTCACTGGTCAACTCCAGCAC |
| *Adipoq* | Mouse | F' TGTTCCTCTTAATCCTGCCCA<br>R' CCAACCTGCACAAGTTCCCTT |
| *Atgl(Pnpla2)* | Mouse | F' ATGTTCCCGAGGGAGACCAA<br>R' GAGGCTCCGTAGATGTGAGTG |
| *Mgl* | Mouse | F' CGGACTTCCAAGTTTTTGTCAGA<br>R' GCAGCCACTAGGATGGAGATG |
| *Il-6* | Mouse | F' TCTATACCACTTCACAAGTCGGA<br>R' GAATTGCCATTGCACAACTCTTT |
| *Il-1b* | Mouse | F' GAAATGCCACCTTTTGACAGTG<br>R' TGGATGCTCTCATCAGGACAG |
| *Ccl-2* | Mouse | F' TAAAAACCTGGATCGGAACCAAA<br>R' GCTGCTTTGCCTACCTCTCC |
| *Ccl-5* | Mouse | F' TCGAGTGACAAACACGACTGC<br>R' GCATTAGCTTCAGATTTACGGGT |
| *Pnpla3* | Mouse | F' TTCTCTGGCCTAATCCCTCCT<br>R' TGACACCGTGATGGTGGTTTT |
| *Lal* | Mouse | F' CTGGTGAGGAACACTCGGTC<br>R' AGCCGTGCTGAAGATACACAA |
| *HSL(LIPE)* | Human | F' TCAGTGTCTAGGTCAGACTGG<br>R' AGGCTTCTGTTGGGTATTGGA |
| *ATGL(PNPLA2)* | Human | F' GAGATGTGCAAGCAGGGATAC<br>R' CTGCGAGTAATCCTCCGCT |
| *MGL* | Human | F' TCGTCAGGGATGTGTTGCAG<br>R' AGGCGAAATGAGTACCATGCC |
| *IL6* | Human | F' ACTCACCTCTTCAGAACGAATTG<br>R' CCATCTTTGGAAGGTTCAGGTTG |
| *CCL2* | Human | F' CAGCCAGATGCAATCAATGCC<br>R' TGGAATCCTGAACCCACTTCT |
| *CCL5* | Human | F' CCAGCAGTCGTCTTTGTCAC<br>R' CTCTGGGTTGGCACACACTT |

| Primers for genotyping PCR | Sequence (5' -> 3') |
|---|---|
| *Fgf21*$^{flox}$-F | CAGGAGAAACAGCCATTCACTTTG |
| *Fgf21*$^{flox}$-R | TAGATCTCATCCATTCCATCAGGG |
| *Adipoq* Cre-F | GAACGCACTGATTTCGACCA |
| *Adipoq* Cre-R | GCTAACCAGCGTTTTCGTTC |
