## Supplementary figures and images for "Cancer-associated adipocytes mediate CD8^+^T cell dysfunction via FGF21-driven lipolysis"

### Graphical Abstract

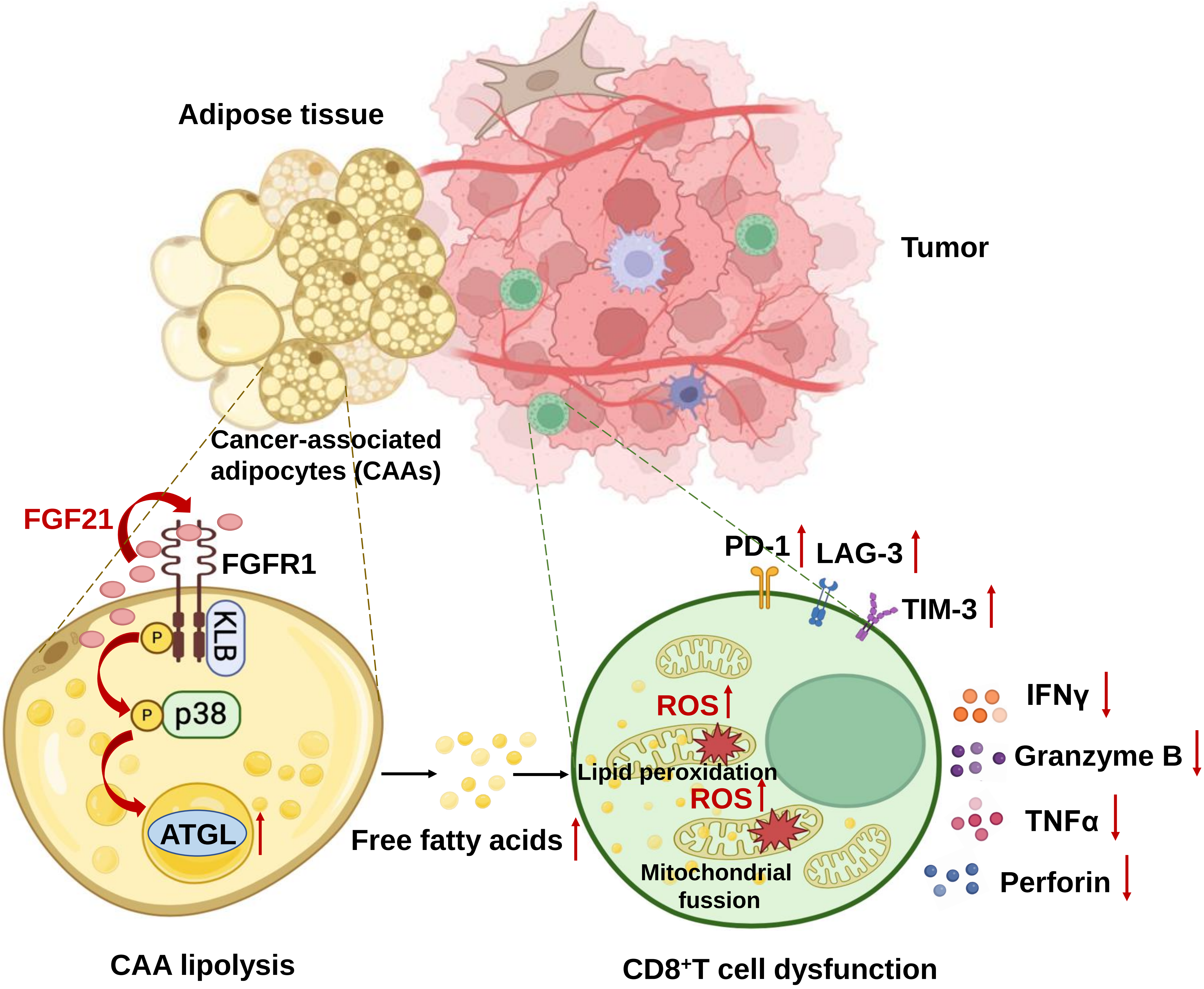
